# A Measurable Systemic Immune Phenotype Links Circulating Myeloid Dysfunction to Tumour Spatial Organization in Gastroesophageal Adenocarcinoma

**DOI:** 10.64898/2026.09.17.752366

**Authors:** Iqraa Dhoparee-Doomah, Yichen Li, Manal Al Dow, Shayan Hajhashemi, Ariane Brassard, Nissim Benizri, Sabrina Leo, Katy Milne, Ming Zi Xu, Qian Qiu, Brad H Nelson, Pierre-Olivier Fiset, Betty Giannias, France Bourdeau, Jonathan Spicer, Lorenzo Ferri, Jean Tchervenkov, Kim Ma, Jonathan Cools-Lartigue

## Abstract

**Background:** Both static and longitudinal measures of circulating neutrophil-to-lymphocyte ratio (cNLR) are prognostic markers across solid malignancies, including gastroesophageal adenocarcinoma (GEA). Despite this consistency, cNLR remains poorly integrated into clinical decision-making and is generally regarded as a nonspecific inflammatory or disease-burden signal rather than a defined biological state. Analysis of a large, prospectively assembled GEA cohort showed that cNLR was only weakly associated with tumour burden or pathological response, yet both baseline and postoperative cNLR independently predicted outcome.

**Methods:** These observations prompted a retrospective biological investigation of what host biology cNLR captures. Using available clinically linked biospecimens, we performed multiscale analyses across complementary patient subsets encompassing peripheral-blood immune function and soluble signalling, tumour single-cell transcriptional programmes, and spatial myeloid organization within tissue.

**Results:** High baseline cNLR was associated with reduced circulating cytotoxic and immune-trafficking mediators, elevated angiogenesis-associated factors, neutrophil-mediated suppression of lymphocyte proliferation, impaired tumour-cell killing, and enhanced spontaneous neutrophil extracellular trap formation. Single-cell profiling identified coordinated transcriptional differences across tumour compartments, including innate, cytokine and myeloid programmes. Spatial profiling showed that myeloid abundance and organization varied across anatomical compartments; greater tumour-periphery PD-L1⁺ myeloid clustering was associated with inferior recurrence-free survival. Baseline cNLR showed little relationship with static myeloid abundance, whereas pretreatment cNLR dynamics were associated with altered cross-compartment CD8⁺– myeloid spatial relationships. Notably, higher early postoperative cNLR was associated with greater PD-L1⁺ myeloid clustering at the tumour periphery, linking the spatial architecture of the resected tumour with a systemic myeloid phenotype persisting after surgery

**Conclusions:** The data identify a prognostic, multilevel myeloid-associated host phenotype in GEA that is incompletely explained by tumour burden. Integrating systemic immune function, tumour single-cell transcriptomics and spatial myeloid organization demonstrates that these biological domains provide distinct, yet convergent views of the host immune phenotype captured by cNLR. These findings establish a biological framework for the clinical utility of cNLR and highlight myeloid-associated immune processes as promising targets for therapeutic intervention.

**SUMMARY BOX:** **What is already known on this topic:** Circulating neutrophil-to-lymphocyte ratio (cNLR) is a robust prognostic marker across solid malignancies, including gastroesophageal adenocarcinoma, but is generally regarded as a nonspecific marker of inflammation or tumour burden.

**What this study adds:** This study identifies elevated cNLR as a marker of a multilevel myeloid-associated host phenotype in GEA, where it captures complementary systemic and tissue-level immune features that are inadequately explained by tumour burden.

**How this study might affect research, practice or policy:** These findings provide a biological framework for interpreting cNLR as an accessible indicator of systemic myeloid-associated immune state. They support prospective evaluation of cNLR as an accessible biomarker for risk stratification and treatment monitoring and highlight myeloid-associated immune processes as potential therapeutic targets.

## 1.0 BACKGROUND

Modern cancer treatment remains predominantly tumour-centered, relying on surgery, radiation, cytotoxic therapy and tumour-intrinsic targeted therapies. Yet patients with comparable histology, stage and molecular features can follow markedly different clinical trajectories, indicating that tumour-centred variables alone do not fully explain disease behaviour (1, 2). The clinical success of immune-checkpoint inhibitors has made this limitation increasingly apparent by establishing host immunity as both a determinant of outcome and a therapeutically actionable component of disease (3–7). In gastroesophageal adenocarcinoma (GEA), immune-checkpoint inhibition improves survival in metastatic disease and reduces recurrence following treatment of localized tumours, further demonstrating the importance of host immunity (3, 5, 8, 9).

The hematopoietic system is central to this interaction. Tumour-and host-derived signals shape hematopoietic production, immune-cell mobilization, circulating soluble mediators and tissue immune organization, generating systemic phenotypes that either constrain or support malignant growth (10–13). Circulating leukocyte counts represent an accessible output of this system and, through repeated routine measurements, capture its dynamics over time.

The circulating neutrophil-to-lymphocyte ratio (cNLR) is one of the simplest composite measures of this output. Elevated cNLR consistently predicts inferior outcomes across solid malignancies, including GEA, often independently of conventional clinicopathological variables (14–17). Despite this reproducibility, cNLR remains poorly integrated into clinical decision-making and is generally interpreted as a nonspecific correlate of inflammation or disease burden. This interpretation does not adequately explain its persistent prognostic association across tumour types, disease stages and treatment settings.

Two biological interpretations are therefore possible. If cNLR is principally a consequence of tumour burden, it should correspond closely with anatomical disease extent and pathological response. Alternatively, cNLR may provide an accessible readout of a broader hematopoietic state rather than simply reflecting independently harmful neutrophils and protective lymphocytes. Under this interpretation, cNLR should retain prognostic information beyond stage and response and correspond to coordinated biological changes across compartments.

Neutrophil plasticity itself provides a plausible example of the latter possibility. Preclinical studies have shown conflicting roles in tumor progression. However, tumour-associated emergency myelopoiesis and neutrophil reprogramming can promote immune suppression, neutrophil extracellular trap formation and metastasis (18). These disparate and widespread effects are in line with the concept that a systemic hematopoietic phenotype may manifest differently across anatomical compartments, with circulating blood, soluble signalling networks and tissue microenvironments providing complementary expressions of a shared underlying host process.

GEA provides a clinically important setting to test this framework, as outcomes remain heterogeneous and frequently poor despite multimodal therapy, including in apparently localized disease (19). Using a large clinically annotated cohort, we first determined whether baseline and longitudinal cNLR measurements primarily reflected anatomical stage or pathological response or provided independent prognostic information. We then examined the biological phenotype associated with cNLR across complementary translational cohorts encompassing peripheral-blood (PB) immune function and soluble signalling, tumour single-cell transcriptional programmes, and spatial myeloid organization across histologically normal and tumour tissue compartments. Through this multiscale analysis, we tested whether cNLR represents a nonspecific correlate of cancer burden or an accessible measurement of a distributed, clinically relevant host immune phenotype.

## 2.0 METHODS

### 2.1 Patient Selection and Samples

The Esophageal and Gastric Data-and Biobank at the McGill University Health Centre (Nagano protocol MP-37-2007-856) included 1,451 patients with histologically confirmed GEA diagnosed between January 2011 and October 2024. Clinical characteristics, treatment information, pathological stage and outcomes were abstracted from clinical records. Complete blood count (CBC) measurements were available for 909 patients, of whom 544 had sufficient temporal information for baseline cNLR measurement. Analysis-specific eligibility yielded 464 patients for clinical-stage analysis, 417 patients for locked baseline-survival analysis, and 189 patients for recovery-window/TRG landmark analysis. Cohort derivation and baseline characteristics are summarized in **Supplementary Figure 1** and **Supplemental Table S1**.

Patients with squamous-cell carcinoma, concomitant active primary malignancy, or clinically documented concurrent infectious or inflammatory conditions at biospecimen collection were excluded from analyses. PB and tumour specimens were obtained through the biobank, with histopathological review performed before tissue-based analysis. Fresh tissue for single-cell RNA sequencing was processed according to procedures described in **section 2.4.1**.

The biobank and associated translational studies were approved by the McGill University Health Centre Research Ethics Board under protocol 2024-9907. All participants provided written informed consent.

### 2.2 Retrospective derivation of cNLR measurements

cNLR was calculated by dividing the absolute neutrophil count by the absolute lymphocyte count obtained from the same CBC. CBCs with a missing or non-positive lymphocyte count were excluded. When multiple CBCs were obtained on the same calendar day, values were collapsed to the daily median.

Temporal cNLR definitions were specified retrospectively (**Fig. 1A**). Baseline and pretreatment windows were designed to characterize the circulating inflammatory state before cancer-directed therapy, whereas postoperative measures were selected to characterize recovery beyond the immediate postoperative period, consistent with prior studies evaluating persistent NLR several months after gastrectomy (20–22).

**Figure 1.**
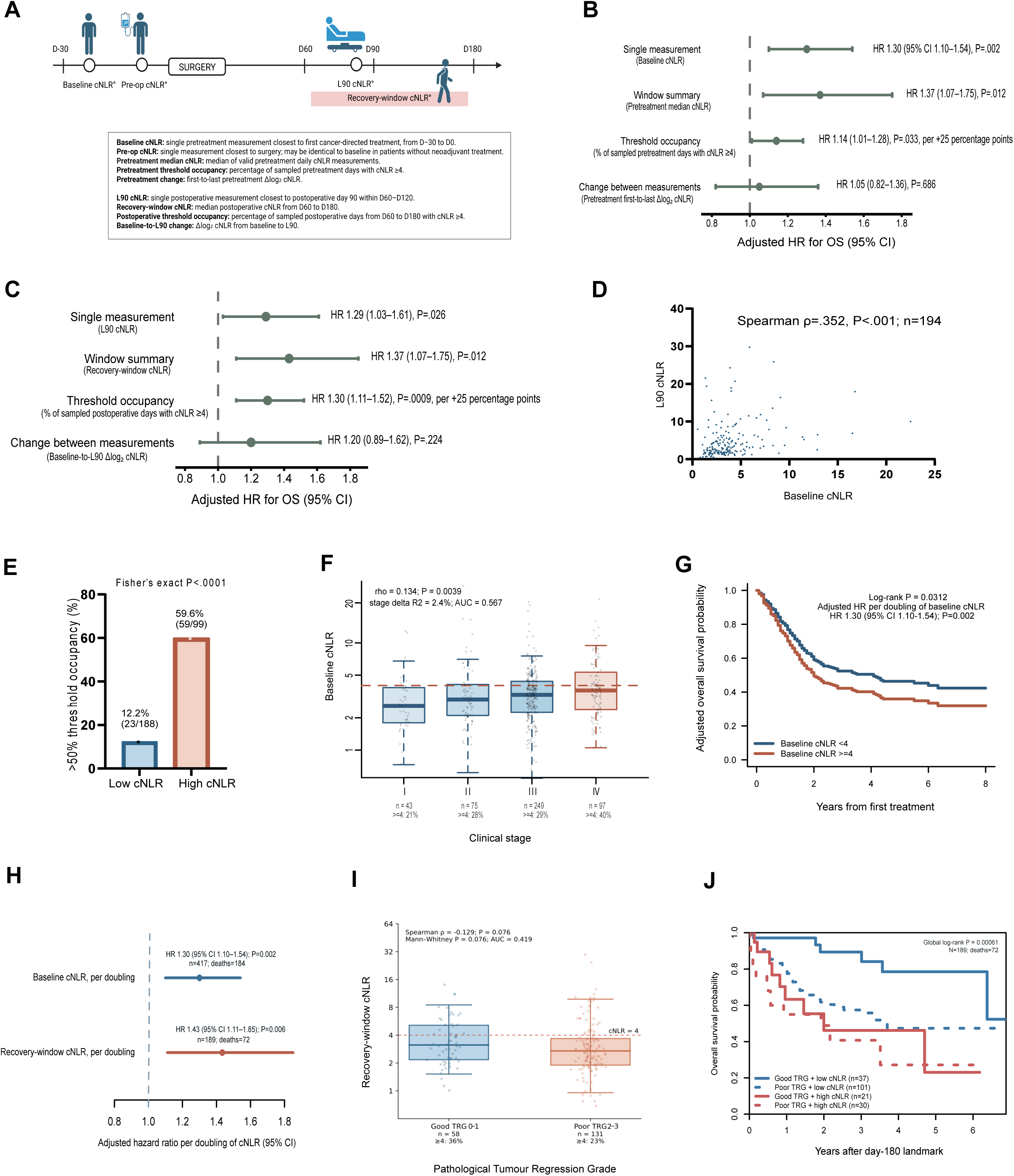
cNLR provides prognostic information beyond tumour stage and pathological response. (A) Timing and definitions of baseline, preoperative, pretreatment, L90 and recovery-window cNLR measurements. (B-C) Adjusted associations of (B) pretreatment and (C) postoperative cNLR summaries with OS, including single measurements, window summaries, threshold occupancy and longitudinal change. Hazard ratios for single cNLR measurements and window summaries are expressed per doubling; threshold-occupancy HRs are expressed per 25-percentage-point increase. (D) Relationship between baseline and L90 cNLR among patients with paired measurements. (E) Proportion of patients with >50% pretreatment threshold occupancy according to baseline cNLR category. (F) Baseline cNLR according to clinical stage (G) OS according to baseline cNLR in the locked 417-patient survival cohort. (H) Adjusted HRs for OS per doubling of baseline and recovery-window cNLR. (I) Recovery-window cNLR according to pathological tumour regression grade (TRG). (J) Day-180 landmark OS according to combined TRG and recovery-window cNLR groups. Survival curves were compared using log-rank tests, and HRs are shown with 95% CI. **Abbreviations:** cNLR: circulating neutrophil-to-lymphocyte ratio; OS: overall survival; HRs: Hazard ratio; TRG: tumour regression grade

The date of first cancer-directed treatment defined day 0. Baseline cNLR was the single valid cNLR measurement closest to first cancer-directed treatment within day −30 to 0. Preoperative cNLR was the single valid measurement closest to surgery and could be identical to baseline cNLR in patients undergoing surgery without neoadjuvant treatment. Pretreatment longitudinal summaries were derived from valid daily pretreatment cNLR measurements. Pretreatment median cNLR was the median of available daily pretreatment measurements. Pretreatment threshold occupancy was the proportion of sampled pretreatment days with cNLR ≥4. Pretreatment change was expressed as the first-to-last pretreatment Δlog₂ cNLR, calculated as log₂(last pretreatment cNLR / first pretreatment cNLR). L90 cNLR was the single valid postoperative measurement closest to postoperative day 90 within days 60 to 120 (22). Recovery-window cNLR was the median of valid postoperative daily cNLR measurements from days 60 to 180. Postoperative threshold occupancy was the proportion of sampled postoperative days from days 60 to 180 with cNLR ≥4. Baseline-to-L90 change was defined as Δlog₂ cNLR, calculated as log₂(L90 cNLR / baseline cNLR). Threshold occupancy represents the proportion of observed CBC days meeting the prespecified cNLR threshold, reflecting persistence of an elevated inflammatory phenotype across serial measurements rather than continuous time above threshold (23).

For analyses requiring categorical stratification, cNLR <4 was classified as low and cNLR ≥4 as high, using an a priori threshold supported by prior GEA studies (17, 24, 25). Continuous cNLR values were retained where appropriate and log₂-transformed in Cox models so that hazard ratios represented the effect of a doubling in cNLR. Threshold-occupancy effects were expressed per 25-percentage-point increase where indicated. Additional details are provided in **Supplementary Methods S1.1.1.**

### 2.3 Peripheral-blood immune phenotype associated with high cNLR

Pretreatment PB was collected before initiation of cancer-directed therapy, thereby limiting the influence of heterogeneous treatments on immune composition and functions. PB mononuclear cells (PBMCs), neutrophils and plasma were isolated from the same blood draw when sample volume and processing permitted. Detailed translational cohort derivation is included in **Supplementary Figure S1**. Assay-specific subsets were determined by biospecimen availability, cell yield, viability and experimental requirements.

Each patient constituted one biological replicate; technical replicates were averaged before statistical analysis unless otherwise specified.

#### 2.3.1 Isolation of PBMCs and neutrophils from peripheral blood

PB was processed promptly after collection, with PBMCs isolated by density-gradient centrifugation, and neutrophils recovered from the granulocyte fraction using dextran sedimentation and red blood lysis. Cell number and viability were assessed, and only samples meeting prespecified quality criteria were included. Plasma was separated, aliquoted and stored at −80°C until analysis. Detailed procedures are provided in **Supplementary Methods S1.2.**

#### 2.3.2 PBMC-Neutrophil Co-Culture Assay

PBMCs were cultured alone or with autologous neutrophils to assess neutrophil-mediated effects on PBMC expansion. PBMCs (1 × 10^6^) were stimulated with anti-CD3 and anti-CD28, with or without 5 × 10^5^ neutrophils, corresponding to a PBMC-to-neutrophil ratio of 2:1. After 5 days, viable PBMCs were quantified using a Vi-CELL automated cell counter. Three technical replicates were averaged per condition, and relative viable PBMC count was calculated as the mean PBMC count in coculture divided by the mean count in matched PBMC-alone wells. Values below 1 indicated reduced viable PBMC expansion in the presence of neutrophils. Detailed experimental procedures are provided in **Supplementary Methods S1.3**.

#### 2.3.3 PBMC-Mediated Cytotoxicity Assay

PBMC-mediated antitumour activity was assessed using A549-GFP cells as a standardized adherent target. A549-GFP cells were cocultured for 24 hours with PBMCs recovered from the preceding functional assay at an effector-to-target ratio of 50:1. Residual adherent GFP-positive tumour-cell area was quantified from one central image per well using ImageJ v2.1, with lower GFP-positive area indicating greater PBMC-mediated cytotoxic activity. Six technical replicates were averaged to generate one intra-patient value per condition. Image acquisition and quantification were performed blinded to cNLR group. Comprehensive experimental procedures are provided in **Supplementary Methods S1.4**.

#### 2.3.4 NET Formation and Staining

Spontaneous NET formation was assessed in freshly isolated neutrophils incubated without stimulation for 4 hours, with PMA-treated cells included as positive controls. NETs were visualized using SYTOX Green under identical imaging conditions. NET formation was confirmed by immunofluorescence staining for citrullinated histone H3 (H3Cit) and CD66b. Four randomly selected fields were quantified per condition using ImageJ v2.1, with H3Cit-positive area normalized to CD66b-positive area for each field. Image acquisition and quantification were blinded to cNLR group. Detailed experimental procedures are provided in **Supplementary Methods S1.5**.

#### 2.3.5 Multiplex Cytokine and Chemokine Profiling of Patient Plasma

Baseline plasma from treatment-naïve GEA patients were analysed using the Human Cytokine/Chemokine 96-Plex Discovery Assay (EVE Technologies). Samples were run in duplicate and averaged to generate a single analyte value per patient.

For predefined biological signatures, analyte concentrations were standardized as z-scores and averaged with equal weighting to generate composite scores. Patients with missing values for any component analyte were excluded from that composite analysis.

Detailed analyte lists, composite-score definitions, and statistical procedures are provided in **Supplementary Methods S1.6** and **Supplementary Table S2**.

### 2.4 Tumour transcriptional programmes associated with high cNLR

A single-cell RNA-sequencing resource was available from 22 GEA patients (n = 13 low-,9 high-cNLR patients) and was used for whole-tumour and cell-compartment transcriptional analyses. Neutrophil-specific analyses were restricted to 7 pre-treatment tumour specimens with sufficient neutrophil representation (n=3 low cNLR; n=4 high cNLR). These samples were used to determine whether low-and high-cNLR states were associated with distinct intratumoural transcriptional programmes. Patient-level pseudobulk analyses were used for differential-expression and pathway-level comparisons to avoid treating individual cells as independent observations.

#### 2.4.1 Single-cell RNA Sequencing and Data Processing

Fresh tumour tissue was enzymatically dissociated to generate single-cell suspensions. Samples meeting predefined quality thresholds for viability, erythrocyte contamination, and debris were processed using the 10x Genomics Chromium platform with Next GEM Single Cell 3′ v3.1 chemistry. Libraries were sequenced on DNBSEQ-G400, NovaSeq 6000, or HiSeq 4000 instruments.

Sequencing reads were quantified against the GRCh38 transcriptome using kallisto (0.46.2) and bustools (0.40.0). Cells with fewer than 1,000 UMIs, fewer than 500 detected genes, or greater than 10% mitochondrial reads were excluded. Putative doublets were removed using Scrublet (0.2.3) (26). Following quality control, a total of 73,844 cells from 26 samples were retained for downstream analyses. Data were normalized, log-transformed, and integrated using Harmony (1.2.4) following selection of 4,000 highly variable genes. Graph-based clustering was performed using the Leiden algorithm, and cellular distributions were visualized using uniform manifold approximation and projection.

#### 2.4.2 Cell Annotation and Transcriptional State Analysis

Broad cell identities were assigned by reference mapping to the Human Cell Landscape and subsequently refined using canonical marker expression (27–29). Exploratory differential expression of pre-selected genes between cNLR groups was assessed at the single-cell level in neutrophils and T cells using the Wilcoxon rank-sum test implemented in Seurat (5.4.0). Functional states within neutrophil compartment were quantified using curated gene signatures and UCell scores were assessed at single-cell level (30–35). For pathway-level analyses, expression was aggregated within each sample and cell type to generate pseudobulk profiles; Reactome and Hallmark pathway activity were then assessed by GSVA version 2.4.4, with scores averaged across multiple samples from the same patient to obtain one patient-level value (36, 37).

Detailed experimental procedures are provided in **Supplementary Methods S1.7 and S1.8.**

### 2.5 Tissue myeloid organization associated with the systemic cNLR phenotype

#### 2.5.1 Archival Tissue Microarray Cohort

Spatial tissue analyses used an archival GEA tissue microarray constructed in 2016, independently of this study and without cNLR-based selection. The array included tissue from 39 clinically annotated patients and, where available, four anatomical compartments: histologically normal esophagus and stomach, primary tumour core, and primary tumour periphery. Tissue availability differed across patients and compartments, resulting in analysis-specific numbers of evaluable patients and cores. Patient characteristics are provided in **Supplemental Table S1**.

Representative regions were selected by a pathologist following review of haematoxylin and eosin–stained sections. Tumour core was defined as central malignant tissue, tumour periphery as the tumour–stroma interface, and normal esophageal and gastric samples as histologically non-malignant mucosa without tumour involvement.

Clinical records and available CBC measurements were subsequently linked to the archival tissue cohort. Blood-based analyses were therefore retrospective and limited to patients with assignable cNLR measurements within the predefined temporal windows. The TMA was not enriched or balanced according to cNLR category, clinical outcome or tissue phenotype.

#### 2.5.2 Multiplex Immunostaining

Chromogenic immunohistochemistry for CD3 and neutrophil elastase was performed to derive a tissue neutrophil-to-lymphocyte ratio. Multiplex immunofluorescence was subsequently performed using the Akoya OPAL tyramide signal-amplification system on an IntelliPATH FLX platform, with CD66b, CD4, CD8, FOXP3, CD68, PD-L1, pan-cytokeratin and DAPI. Single marker, autofluorescence and tissue controls were used for assay optimization, validation and spectral unmixing.

Slides were imaged at 20× magnification using the Vectra Polaris system (Akoya Biosciences). Detailed experimental procedures are provided in **Supplementary Methods S1.9** and **Supplementary Tables S3.**

#### 2.5.3 Image segmentation and cellular phenotyping

Images were analysed using inForm v10.1.2. A supervised machine-learning classifier trained on 15 representative fields was used to segment epithelial and stromal tissue regions and identify individual cells. Eleven mutually exclusive cellular phenotypes were assigned using predefined marker-expression rules, while preserving the X–Y coordinates and tissue-compartment designation of each cell. Phenotype definitions and quality-control criteria are provided in **Supplementary Methods S1.10** and **Supplementary Table S4**.

PhenoptrReports v0.3.3 (Akoya Biosciences) was used to generate patient-and core-level cell counts, densities and spatial outputs within epithelial and stromal compartments. Core-level measurements were subsequently summarized at the patient level according to the prespecified analysis plan.

### 2.6 Spatial Analyses

Spatial analyses were performed using single-cell phenotypes and X–Y centroid coordinates. Abundance was summarized as phenotype-positive cell counts and densities per mm² of analysed tissue. Cellular relationships were quantified using centre-to-centre nearest-neighbour distances and the proportions of CD8⁺ T cells within prespecified 25-, 50-, and 100-µm radii of myeloid population (38). Within-population organization was assessed by comparing observed nearest-neighbour distances with those expected under complete spatial randomness, following the general Clark–Evans nearest-neighbour framework (39); positive log₂ scores indicated shorter-than-expected intercellular spacing and greater clustering.

Core-level measurements were aggregated to patient-level summaries within each anatomical compartment. Premalignant-field tissue was defined as Barrett’s esophagus, Barrett’s esophagus with dysplasia, or dysplasia. Cross-compartment gradients were calculated from patient-level compartment summaries. Tumour CD66b⁺ abundance was summarized across tumour core and tumour periphery.

Patient-level tissue features were related to baseline cNLR, postoperative cNLR, and ΔcNLR (postoperative cNLR − baseline cNLR) using Spearman correlation. cNLR group comparisons used the prespecified threshold of <4 versus ≥4; other spatial measures were dichotomized at cohort median where indicated. For visualization of continuous clustering–RFS association, model-estimated survival curves were displayed at the first and third quartiles of the clustering score. Associations with survival or recurrence were evaluated using Kaplan–Meier analysis, log-rank testing, and Cox proportional-hazards models. Outcome definitions, censoring and eligibility criteria are provided in **Supplementary Methods S1.11.5,** with full analysis workflows in **Supplementary Methods S1.11.**

### 2.7 General Statistical Analysis and NLR Definitions

Continuous variables were summarized as median and interquartile range and categorical variables as number and percentage. Comparisons between independent groups were performed using the Mann–Whitney U test or Welch’s t test, as appropriate. Paired comparisons were performed using the Wilcoxon signed-rank test, and associations between continuous variables were assessed using Spearman correlation. Technical replicates were averaged to generate one value per patient before inferential analysis unless otherwise specified.

Survival outcomes were analysed using Kaplan–Meier methods, log-rank tests and Cox proportional-hazards regression. For baseline cNLR analyses, OS was measured from first cancer-directed treatment. The locked complete-case baseline model included clinical stage, age, sex, clinical cohort and log₂-transformed baseline cNLR; the analysis comprised 417 patients and 184 deaths. For postoperative analyses, OS was evaluated using a day-180 landmark among patients with evaluable recovery-window cNLR and TRG. Recovery-window cNLR was analysed both categorically (<4 versus ≥4) and continuously on the log₂ scale; continuous models included TRG and were stratified by primary tumour site. For the alternative cNLR summaries shown in **Fig. 1B-C**, hazard ratios for single measurements and window summaries were expressed per doubling of cNLR, threshold-occupancy effects per 25-percentage-point increase, and longitudinal-change effects per unit Δlog₂ cNLR. Proportional-hazards assumptions were assessed for Cox models. All statistical tests were two-sided, and P values below .05 were considered statistically significant. Where multiple related features were evaluated, false-discovery rates were controlled using the Benjamini–Hochberg procedure. Analyses were performed using GraphPad Prism v10.6.1 and R version 4.5.1.

Assay-specific statistical procedures are described in the corresponding **Methods sections** and **Supplementary Methods**.

## 3.0 RESULTS

### 3.1 Clinical observation: NLR provides prognostic information beyond anatomical disease extent and pathological response

Routine CBC measurements were aligned to treatment and surgery dates to derive complementary single-measurement, window-summary, threshold-occupancy and longitudinal-change measures of cNLR (**Fig. 1A**). Prognostic associations depended on how and when cNLR was summarized. Baseline cNLR (adjusted HR per doubling 1.30, 95% CI 1.10–1.54; P=.002), pretreatment median cNLR (HR 1.37, 95% CI 1.07–1.75; P=.012) and pretreatment threshold occupancy (HR 1.14 per 25-percentage-point increase, 95% CI 1.01–1.28; P=.033) were associated with mortality, whereas first-to-last pretreatment change was not (HR 1.05, 95% CI 0.82–1.36; P=.686; **Fig. 1B**). Postoperatively, L90 cNLR (HR 1.29, 95% CI 1.03–1.61; P=.026), recovery-window cNLR (HR 1.37, 95% CI 1.07–1.75; P=.012) and postoperative threshold occupancy (HR 1.30 per 25-percentage-point increase, 95% CI 1.11–1.52; P=.0009) were prognostic, whereas baseline-to-L90 change was not (HR 1.20, 95% CI 0.89–1.62; P=.224; **Fig.1C**).

Baseline cNLR only partially reflected the subsequent postoperative state. Among 194 patients with paired baseline and L90 measurements, the two were positively but incompletely correlated (Spearman ρ=.352, P<.001; **Fig. 1D**). High baseline cNLR also identified a more persistent pretreatment phenotype: >50% threshold occupancy occurred in 59.6% of patients with baseline cNLR ≥4 compared with 12.2% of those with baseline cNLR <4 (59/99 versus 23/188; Fisher’s exact P<.0001; **Fig. 1E**).

Despite this limited relationship with anatomical disease extent (**Fig. 1F**), baseline cNLR remained prognostically informative. In the locked 417-patient survival cohort, high baseline cNLR was associated with inferior OS (n= 283 low-, 134 high-cNLR patients; log-rank P=.0312; **Fig. 1G**). After adjustment for clinical stage, age, sex and cohort, each doubling of baseline cNLR remained independently associated with mortality (adjusted HR 1.30, 95% CI 1.10–1.54; P=.002; 184 deaths; **Fig. 1G-H; Supplementary Table S5**). We next examined whether postoperative cNLR reflected pathological treatment response. Recovery-window cNLR was not significantly associated with pathological tumour regression grade (TRG) (n = 189 patients; Spearman ρ=−.129, P=.076; Mann– Whitney P=.076; AUC=.419; **Fig. 1I**). Nevertheless, TRG and recovery-window cNLR provided complementary prognostic information: the four groups defined jointly by TRG and cNLR showed significantly different OS in the day-180 landmark analysis (global log-rank P=.00061; **Fig. 1J**). In the mutually adjusted categorical model, both poor TRG (HR 2.14, 95% CI 1.20–3.82; P=.010) and recovery-window cNLR ≥4 (HR 2.57, 95% CI 1.53– 4.29; P<.001) were independently associated with mortality (**Supplementary Table S6**). When analysed continuously, each doubling of recovery-window cNLR was associated with increased mortality (adjusted HR 1.43, 95% CI 1.11–1.85; P=.006; **Fig. 1H; Supplementary Table S6**).

These findings indicate that cNLR is only modestly related to clinical stage and shows little relationship with pathological response yet remains prognostic before treatment and during postoperative recovery. This motivated subsequent investigation of the host immune phenotype represented by elevated cNLR.

### 3.2 Peripheral-blood immune phenotype associated with high cNLR

To minimize the influence of treatment-related immune perturbation, all PB analyses used blood collected before initiation of cancer-directed therapy. We first assessed whether high baseline cNLR was accompanied by broader changes in circulating soluble mediators and then whether it corresponded to functional differences in neutrophils and lymphocytes.

#### 3.2.1 High cNLR Identifies a Distinct Soluble Immune Microenvironment

Plasma from 31 treatment-naïve patients (17 low cNLR; 14 high cNLR) was profiled for 96 soluble immune and inflammatory mediators (**Fig. 2A**). High cNLR patients showed reduced circulating granzyme A (p = 0.0483; **Fig. 2B**) and perforin (p = 0.0060; **Fig. 2B**). A composite cell-death score incorporating soluble Fas, granzyme A, granzyme B, perforin and TRAIL was also reduced (p = 0.0224; **Fig. 2C**), consistent with attenuated systemic cytotoxic signalling.

**Figure 2.**
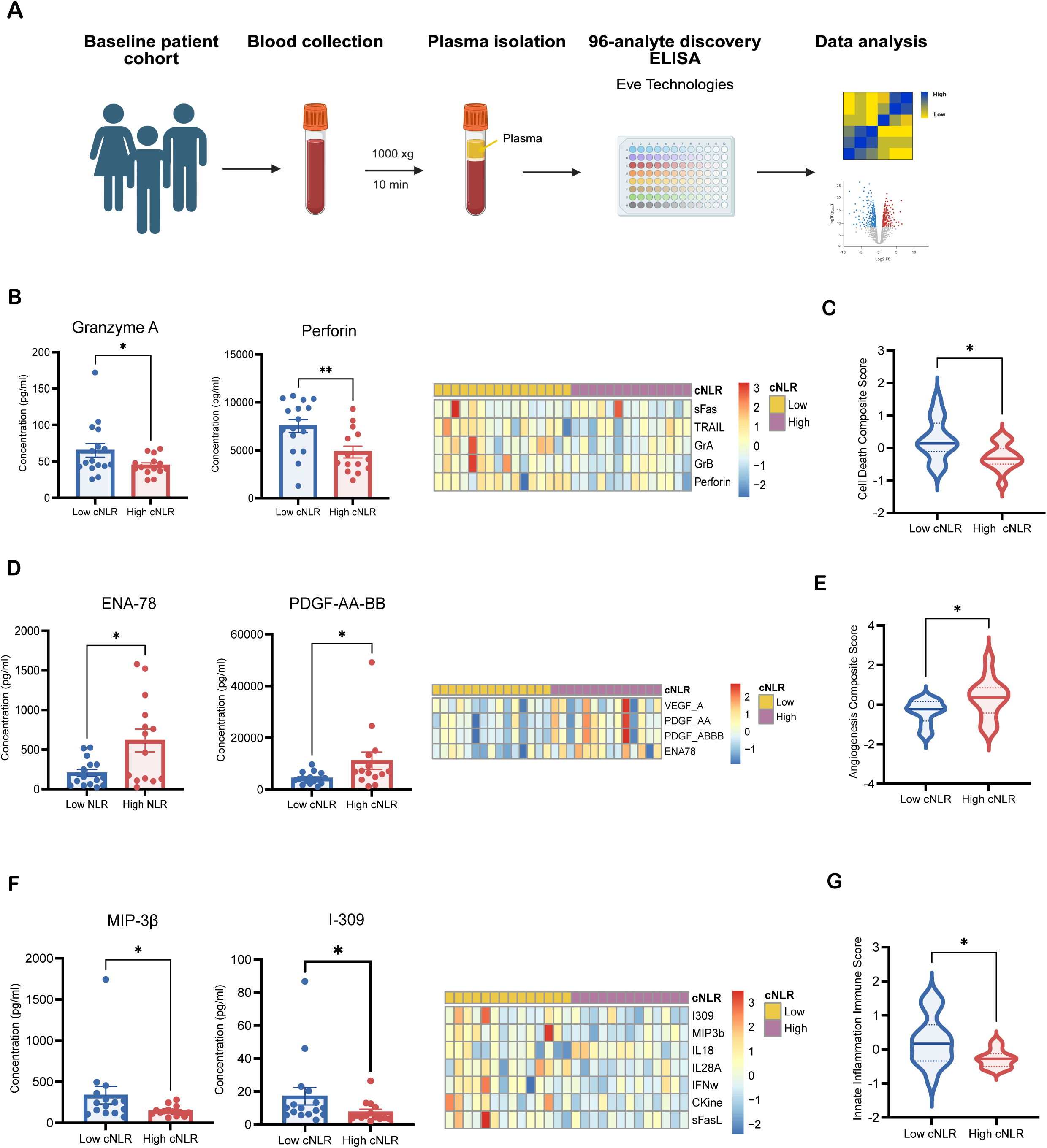
High baseline cNLR defines a distinct systemic soluble immune environment. (A) Plasma was isolated from baseline blood samples obtained from treatment-naïve gastroesophageal adenocarcinoma patients and analyzed using a 96-analyte discovery ELISA panel. Patients were stratified according to low cNLR (n=17) or high cNLR (n=14). (B) Concentrations of selected cell-death-associated mediators and heatmap of cell-death markers across individual patients. (C) Cell-death composite score according to cNLR group. (D) Concentrations of selected angiogenesis-associated mediators and heatmap of angiogenesis markers. (E) Angiogenesis composite score according to cNLR group. (F) Concentrations of selected immune-signalling and trafficking mediators and corresponding patient-level heatmap. (G) Immune-trafficking composite score according to cNLR group. Heatmaps display row-scaled analyte concentrations, with the upper annotation bar indicating cNLR group. Composite scores were calculated using a z-score-based integration approach, where individual analytes were standardized across the cohort and averaged to generate a composite biological signature score. Between-group comparisons were performed using two-tailed Mann– Whitney tests. *P < .05; **P < .01. **Abbreviations:** cNLR, circulating neutrophil-to-lymphocyte ratio; CCL5, C-C motif chemokine ligand 5; CKine, chemokine; ELISA, enzyme-linked immunosorbent assay; ENA-78, epithelial neutrophil-activating peptide 78; GEA, gastroesophageal adenocarcinoma; GrA, granzyme A; GrB, granzyme B; IFN-γ, interferon gamma; IL, interleukin; MIP-3β, macrophage inflammatory protein 3 beta; PDGF, platelet-derived growth factor; sFas, soluble Fas; sFasL, soluble Fas ligand; TRAIL, tumour necrosis factor-related apoptosis-inducing ligand; VEGF-A, vascular endothelial growth factor A.

In contrast, ENA-78/CXCL5 (p = 0.0367; **Fig. 2D**) and PDGF-AB/BB (p = 0.0259; **Fig. 2D**) were increased in high cNLR plasma. A composite angiogenesis-associated score comprising of VEGF-A, PDGF-AA, PDGF-AB/BB and ENA-78 was correspondingly elevated in the high-cNLR group (p = 0.0255; **Fig. 2E**). High cNLR was also accompanied by reduced MIP-3β/CCL19 (p = 0.0367; **Fig. 2F**) and I-309/CCL1 (p = 0.0267; **Fig. 2F**); a broader immune-signalling and trafficking composite was lower (P = 0.0255; **Fig. 2G**).

Thus, elevated cNLR did not simply reflect a numerical imbalance between neutrophils and lymphocytes but identified a systemically altered circulating environment characterized by diminished cytotoxic mediators, impaired immune-trafficking signals and enhanced angiogenesis-associated factors.

#### 3.2.2 High cNLR is associated with functional neutrophil reprogramming in GEA

To test whether these soluble differences were accompanied by functional alterations in circulating neutrophils and lymphocytes, we performed autologous PBMC– neutrophil co-culture, PBMC-mediated cytotoxicity, and spontaneous NET assays in nested subsets (**Fig. 3A**).

**Figure 3.**
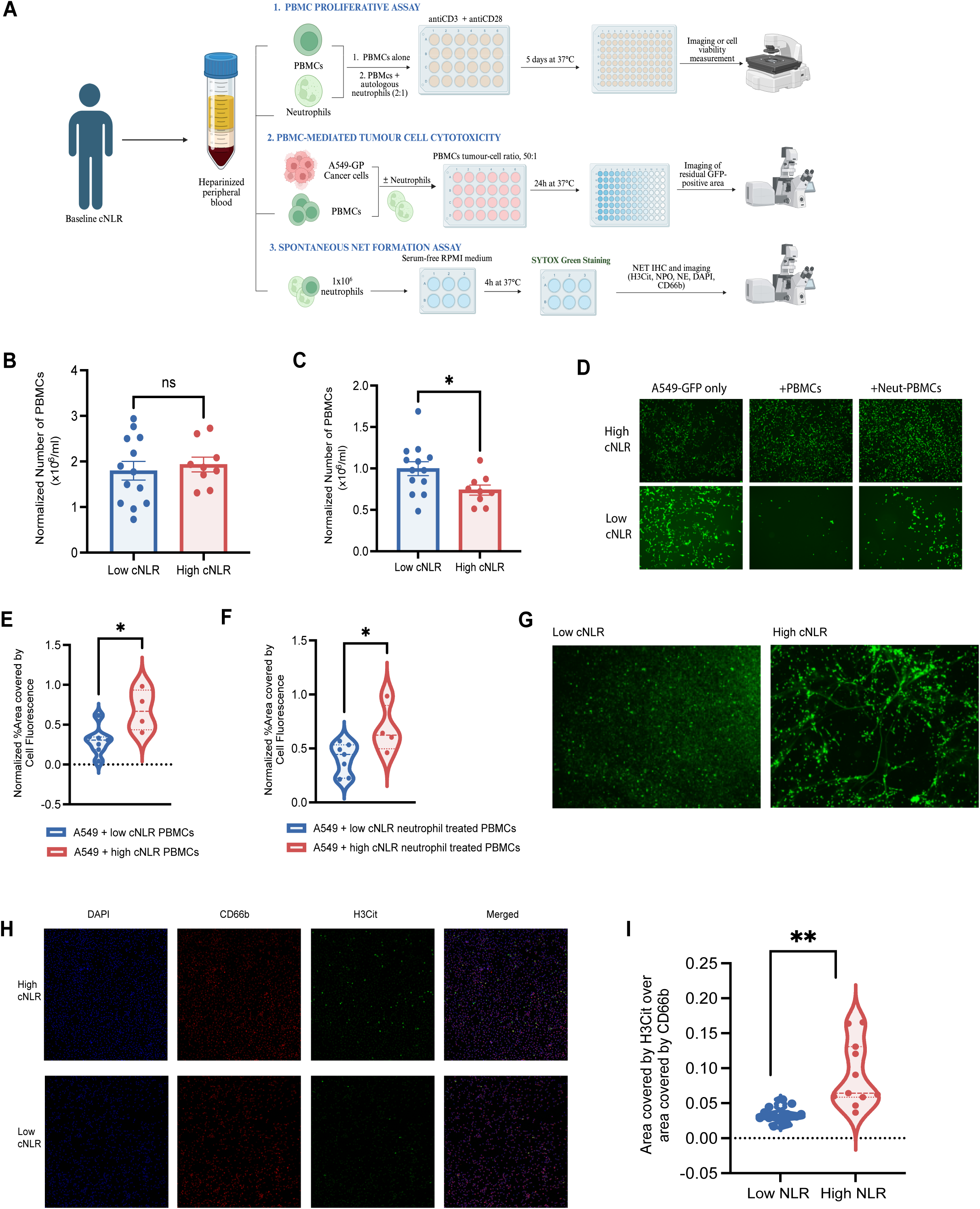
High baseline cNLR neutrophils suppress PBMC proliferation and antitumour activity while promoting NET formation. (A) Experimental overview of PBMC proliferation, A549-GFP cytotoxicity, and spontaneous NET-formation assays. (B) PBMC counts in the absence of neutrophils after 5 days of culture (n=13 low baseline cNLR; 9 high baseline cNLR patients). (C) PBMC counts with autologous neutrophils from low-versus high cNLR patients after 5 days of co-culture (n=13 low baseline cNLR; 9 high baseline cNLR patients). (D) Representative A549-GFP images after co-culture with PBMCs or neutrophil-conditioned PBMCs. (E–F) Quantification of residual A549-GFP area after co-culture with (E) PBMCs or (F) neutrophil-conditioned PBMCs by using ImageJv2.1 (n=7 low baseline cNLR; 4 high baseline cNLR patients). (G) Representative SYTOX Green images of spontaneous NET formation. (H) Representative DAPI, CD66b, H3Cit, and merged immunofluorescence images. (I) H3Cit-positive area normalized to CD66b-positive area. Data represent randomly sampled image fields from 4 low cNLR (16 fields) and 3 high cNLR (11 fields) patients. Linear mixed effects model with cNLR group as a fixed effect and patient as a random effect to account for multiple fields sampled within each patient. Technical replicates were analyzed as described in Methods. Bars show mean ± SEM. ns, not significant; *P < .05; **P < .01. **Abbreviations:** cNLR, circulating neutrophil-to-lymphocyte ratio; GFP, green fluorescent protein; H3Cit, citrullinated histone H3; NET, neutrophil extracellular trap; PBMC, peripheral blood mononuclear cell; SEM, standard error of the mean.

In the absence of neutrophils, viable PBMCs count after five days of stimulation didn’t differ between low- and high-cNLR groups (**Fig. 3B**). When autologous neutrophils were added, PBMC expansion was significantly reduced in high-cNLR patients (**Fig. 3C**). PBMC-mediated cytotoxicity was assessed using A549-GFP cells as a standardized adherent target. Residual GFP-positive area was higher when PBMCs from high-cNLR patients were used, consistent with reduced effector function (**Fig. 3D, E**). Conditioning of PBMCs with autologous high-cNLR neutrophils further increased residual target-cell area (**Fig. 3F**), linking the circulating neutrophil phenotype to impaired lymphocyte-mediated killing of a standardized target.

Finally, neutrophils from high-cNLR patients exhibited greater spontaneous NET formation, confirmed by both SYTOX Green staining and increased citrullinated histone H3 signal normalized to CD66b area (**Fig. 3G–I**).

Taken together, high baseline cNLR identifies a functionally reprogrammed circulating neutrophil state characterized by suppression of lymphocyte proliferation, impaired effector function against a standardized target, and enhanced spontaneous NET formation, accompanied by a soluble milieu depleted of cytotoxic and trafficking mediators and enriched for angiogenesis-associated factors.

### 3.3 Tumour transcriptional programmes associated with high cNLR

To determine whether the pretreatment systemic immune state captured by baseline cNLR was associated with transcriptional differences within the tumour ecosystem, we compared single-cell RNA-sequencing data from 26 evaluable tumour samples from 22 patients (n= 13 low- and 9 high-baseline cNLR).

Analysis of Reactome gene sets showed that low-cNLR tumours were relatively enriched for programmes related to complement activation, antigen processing and presentation and B-receptor signalling (**Fig. 4A**). In contrast, high-cNLR tumours showed relative enrichment of cytokine signalling, innate immune sensing and myeloid-differentiation pathways (**Fig. 4B**).

**Figure 4.**
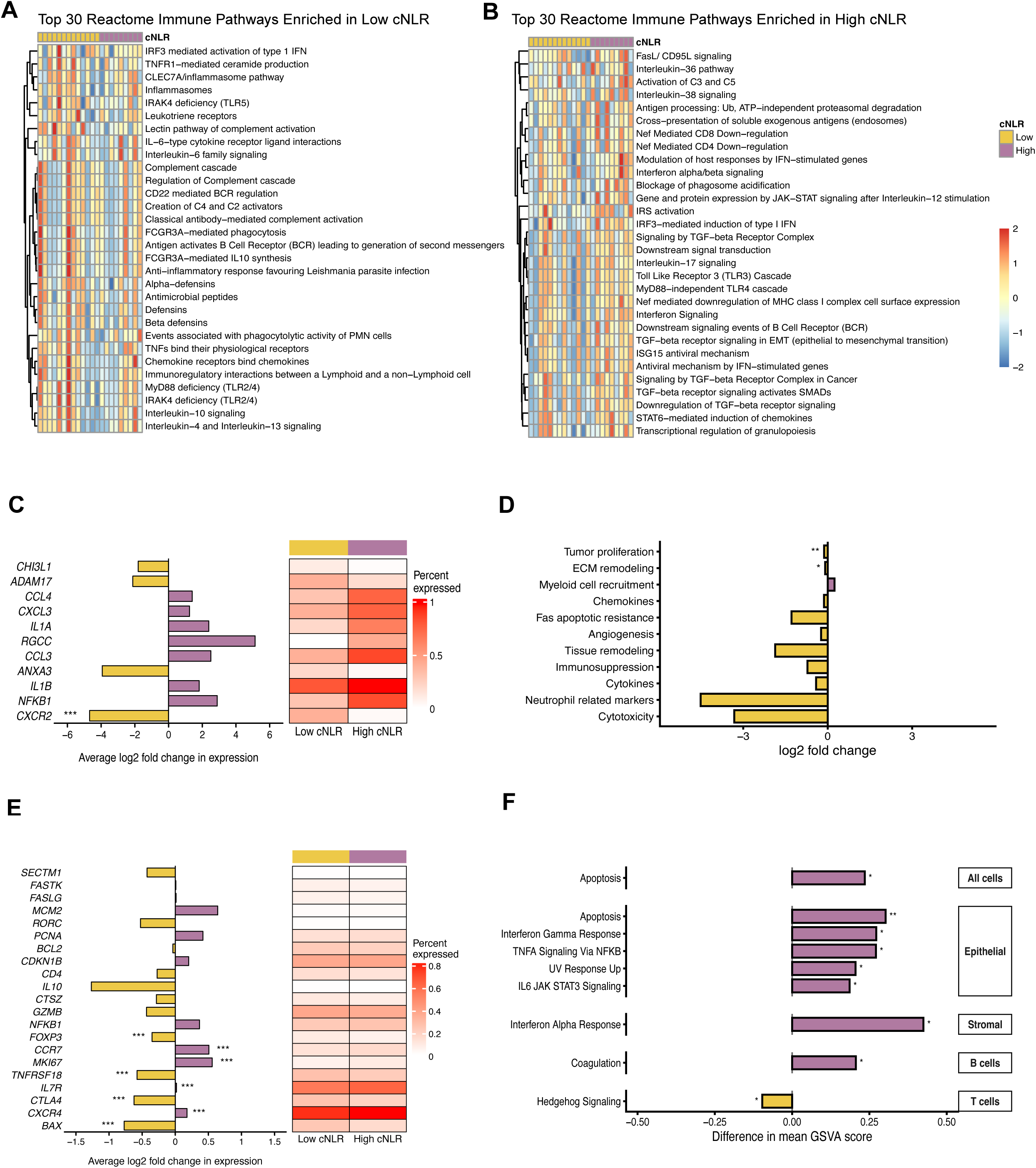
Baseline cNLR is associated with coordinated transcriptional states across different cell compartments. (A–B) Row-scaled GSVA heatmaps of the top Reactome immune pathways relatively enriched in tumours from low-cNLR (n = 13) and high-cNLR (n = 9) patients. (C) Expression of genes of interest in tumour-associated neutrophils assessed at single-cell level. Bars indicate average log₂ fold change. Asterisks indicate adjusted p value (*Padj < .05; **Padj < .01; ***Padj < .001). (D) Selected TAN pathway-enrichment analysis. Asterisks indicate nominal, unadjusted P values (*P < .05; **P < .01; ***P < .001). (E) Expression of genes of interest in tumour-associated T cells assessed at single cell level. Bars indicate average log₂ fold change. Asterisks indicate adjusted p value (*Padj < .05; **Padj < .01; ***Padj < .001). (F) Enriched signalling pathways in different cell compartments in tumours. Enrichment analysis was performed using the HALLMARK gene set database. Asterisks indicate nominal, unadjusted P values (*P < .05; **P < .01; ***P < .001). Bars are coloured by the group with higher expression (yellow, low cNLR; purple, high cNLR). **Abbreviations**: cNLR: circulating neutrophil-to-lymphocyte ratio; GSVA: Gene Set Variation Analysis; TAN; tumour-associated neutrophils

We next examined transcriptional states within the neutrophil compartment. In single-cell level analysis of pre-selected genes, neutrophils from high-cNLR tumours exhibited increased expression of inflammatory and chemokine-associated genes, including *NFKB1, IL1A, IL1B, CCL3, CCL4 and CXCL3*, together with *RGCC* (**Fig. 4C**). Genes implicated in neutrophil trafficking and regulatory function, including CXCR2, *ADAM17* and *ANXA3*, were reduced. Pathway-level analysis further indicated reduced tumour-proliferation and extracellular-matrix-remodelling signatures, with a trend toward increased myeloid-recruitment signalling in high-cNLR tumours (**Fig. 4D**).

Because effective antitumour immunity depends on coordinated interactions between innate and adaptive immune compartments, we next examined tumour-infiltrating T cells. T cells from high-cNLR tumours showed greater expression of genes involved in trafficking and lymph node homing at single-cell level analysis, including *CCR7* and *CXCR4*, together with increased expression of the proliferation marker *MKI67* (**Fig. 4E**). Conversely, low-cNLR T cells showed greater expression of regulatory T cell associated genes, including *FOXP3*, *CTLA4*, and *TNFRSF18*, and the pro-apoptotic gene *BAX* (**Fig. 4E**).

To determine whether baseline cNLR was associated with compartment-specific transcriptional states within the tumour ecosystem, we then used GSVA to analyse Hallmark pathway activity. Overall, high-cNLR tumours have increased apoptotic pathway activity (**Fig. 4F**). A broader transcriptional shift was observed in the epithelial compartment, with nominal significance observed in inflammatory signalling, including TNFα, IFN-γ, and IL6 signalling, together with apoptosis and cellular stress-associated UV-response pathways (**Fig. 4F**). Similarly, the stromal compartment had higher activity in IFN-α response (**Fig. 4F**). B cells from cNLR-high tumours had higher GSVA scores in coagulation, driven predominantly by complement-related genes, while T cells from cNLR-low tumours had higher activity in Hedgehog signalling (**Fig. 4F**). In contrast, no Hallmark pathways differed nominally between cNLR groups in the myeloid compartment. Collectively, these findings show that high baseline cNLR was accompanied by a tumour transcriptional landscape that is not restricted to intratumoral immune cells, suggesting a remodelling throughout the tumour ecosystem.

### 3.4 Tissue myeloid organization associated with the systemic cNLR phenotype

Finally, we examined whether the circulating phenotype associated with cNLR was reflected in tissue myeloid biology. Multiplex spatial profiling of 39 resected GEA specimens quantified myeloid abundance and spatial organization across matched normal, premalignant and tumour compartments (**Fig. 5A**).

**Figure 5.**
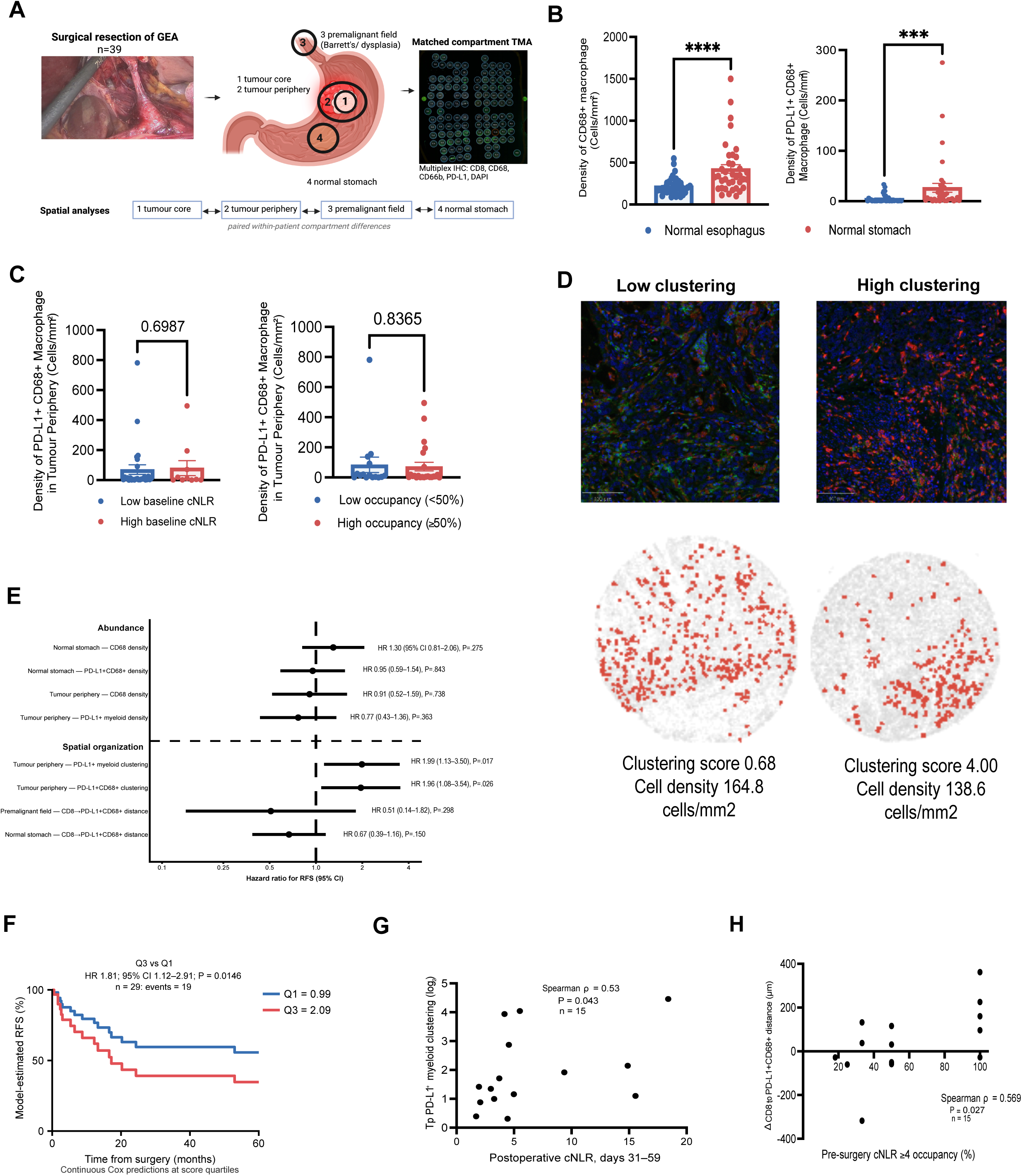
cNLR is associated with compartment-specific differences in myeloid abundance and spatial organization. (A) Matched-compartment spatial profiling strategy in 39 resected GEA specimens. (B) CD68⁺ and PD-L1⁺CD68⁺ macrophage densities in normal esophagus and normal stomach. (C) Tumour-periphery PD-L1⁺CD68⁺ macrophage density according to baseline cNLR and preoperative cNLR threshold occupancy. (D) Representative tumour-periphery regions with contrasting PD-L1⁺CD68⁺ clustering. (E) Associations of compartment-specific myeloid abundance and spatial organization with RFS. (F) Associations of tumour periphery PD-L1+ myeloid clustering quartiles with RFS. (G) Correlation between clustering of PD-L1+ myeloid cells at tumour periphery and post-operative cNLR. (H) Association between preoperative cNLR threshold occupancy and tumour-periphery–premalignant difference in CD8⁺-to-PD-L1⁺CD68⁺ distance. HRs are shown with 95% CIs; survival curves were compared using log-rank tests. Error bars show mean ± SEM. ***P < .001; ****P<.0001. **Abbreviations:** cNLR: circulating neutrophil-to-lymphocyte ratio; GEA: gastroesophageal adenocarcinoma; RFS: Recurrence-free survival; PD-L1: Programmed death ligand 1; Tp: tumour periphery.

Myeloid abundance varied substantially by anatomical compartment. CD68⁺ macrophage and PD-L1⁺CD68⁺ macrophage densities were significantly higher in normal stomach than in normal esophagus (**Fig. 5B**). In contrast, tumour-periphery PD-L1⁺CD68⁺ macrophage density showed little relationship with either baseline cNLR category or preoperative cNLR threshold occupancy (**Fig. 5C**), suggesting that circulating cNLR was not simply a marker of greater local myeloid abundance.

Spatial organization provided additional information beyond cell density. Tumour-periphery regions with similar PD-L1⁺CD68⁺ macrophage densities displayed markedly different clustering patterns (**Fig. 5D**). In continuous survival analyses, greater tumour-periphery PD-L1⁺ myeloid clustering (HR 1.99, 95% CI 1.13–3.50; P=.017) and greater PD-L1⁺CD68⁺ macrophage clustering (HR 1.96, 95% CI 1.08–3.54; P=.026) were associated with inferior RFS, whereas the corresponding abundance measures were not associated with RFS (**Fig. 5E**). Consistent with this, patients in the highest quartile of tumour-periphery PD-L1⁺ myeloid clustering had inferior RFS compared with those in the lowest quartile (HR 1.81; 95% CI 1.12–2.91; P=.0146; n=29; **Fig. 5F**).

In exploratory analyses, we next examined whether these spatial features were related to circulating cNLR. Higher early postoperative cNLR at days 31–59 was associated with greater tumour-periphery PD-L1⁺ myeloid clustering (Spearman ρ=.53, P=.043; n=15; **Fig. 5G**). Preoperative cNLR dynamics also related to cross-compartment immune organization: greater preoperative cNLR threshold occupancy correlated with a larger difference in CD8⁺-to-PD-L1⁺CD68⁺ distance across tissue compartments (Spearman ρ=.569, P=.027; n=15; **Fig. 5H**).

Together, these findings indicate that cNLR relates more closely to myeloid spatial organization than to absolute myeloid abundance. Notably, tumour-periphery PD-L1⁺ myeloid clustering was associated not only with recurrence but also with cNLR measured 31–59 days after surgery, suggesting that spatial features of the resected tumour may be linked to a systemic myeloid phenotype that persists into the postoperative period.

## DISCUSSION and CONCLUSION

Although elevated cNLR predicts outcomes across solid malignancies, its biological meaning remains unclear (14, 40). In our cohort, baseline and recovery cNLR remained prognostic despite weak relationships with stage and pathological response, suggesting that cNLR captures clinically relevant host biology beyond tumour burden.

Translational and sequencing analyses support this interpretation. High cNLR was associated with altered soluble signalling, neutrophil-mediated suppression of PBMC recovery, impaired PBMC-mediated tumour-cell killing, increased spontaneous NET formation and transcriptional differences across multiple cellular compartments. Although these analyses were performed in complementary patient subsets and do not establish intra-patient causality, together they indicate that high cNLR reflects broader immune dysfunction rather than isolated neutrophilia or lymphopenia.

Emergency myelopoiesis provides one plausible biological framework for these findings. Persistent tumour-associated inflammatory signalling can alter myeloid production and mobilization to support tumour progression (11, 12, 41, 42). Our findings are compatible with this process, but we did not directly measure bone-marrow output or immature myeloid populations. cNLR should therefore be viewed as an accessible marker of the resulting systemic immune phenotype rather than as a direct measure of emergency myelopoiesis.

The spatial analyses provide an additional dimension to this phenotype. At the tumour periphery, PD-L1⁺ myeloid clustering, but not myeloid densities, was associated with recurrence. cNLR likewise showed little relationship with static myeloid abundance but was associated with spatial relationships across tissue compartments. Prior studies have shown that spatial immune organization at the tumour-host interface can provide information beyond individual cell abundance (43–45). Our findings thus extend this literature by linking tumour immune architecture to a systemic circulating immune phenotype.

The postoperative findings are particularly informative. Recovery-window cNLR remained prognostic despite being unrelated to pathological response. In a separate exploratory analysis, greater PD-L1⁺ myeloid clustering in the resected tumour was associated with higher cNLR 31–59 days after surgery. Thus, a spatial feature present in the tumour before resection remained related to the circulating immune state after surgery. Although this does not establish causality or exclude occult residual disease, it raises the possibility that tumour-associated immune reprogramming may persist beyond removal of the primary tumour.

This interpretation has implications for how cNLR might be used. The goal would not be to normalize the ratio itself, but to determine whether serial cNLR measurements can identify persistence or re-emergence of an adverse host immune phenotype and whether the underlying biology can be therapeutically modified. Prospective studies incorporating serial blood, marrow and tissue sampling will be needed to determine the mechanisms underlying this phenotype and whether altering them improves clinical outcomes.

Several limitations should be acknowledged. The translational cohorts were assay-specific, sample sizes were modest, and the spatial analyses were exploratory. Longitudinal cNLR definitions were applied retrospectively, and the clinical cohort largely predates routine immune-checkpoint blockade in resectable GEA. Moreover, similar cNLR values may arise through different biological mechanisms. Nevertheless, our findings support cNLR as an accessible marker of a broader systemic immune phenotype that is not fully explained by tumour burden and may persist following tumour resection.

## DECLARATIONS

### Ethics approval and consent to participate

The Esophageal and Gastric Data- and Biobank is registered under Nagano protocol MP-37-2007-856. The biobank and all associated translational research studies were approved by McGill University Health Centre Research Ethics Board (REB protocol 2024-9907). Written informed consent was obtained from all participants.

### Consent for publication

Not applicable. No individual participant data or identifying information are presented in this manuscript.

### Authors’ contributions

IDD performed most of the experimental work, experimental optimization, and associated data analyses, and drafted the manuscript. JCL contributed substantially to the conceptualization of the project. IDD, KM and MAD generated and edited the figures. KM and JCL performed the clinical and spatial analyses and contributed to manuscript review and editing. YL and QQ analyzed the single-cell sequencing data and generated the associated heatmaps and graphical representations. NB, POF, and KM contributed to patient chart review and the compilation and synthesis of clinical data. SH performed the statistical analyses of patient-related clinical data. KMi, BHN, and MZX contributed to the optimization, staining, and initial analysis of the human tissue microarray. AB, SL, JCL, BG, and FB contributed to the experimental work and provided essential resources for completion of this study. MAD, JT, LF, and JS provided constructive feedback on successive drafts of the manuscript. All authors reviewed and approved the final manuscript.

## Acknowledgements

The authors gratefully acknowledge the patients who participated in this study and thank the Thoracic Surgery team at the Montreal General Hospital for their assistance in patient consenting and biospecimen collection. We also acknowledge the Research Institute of McGill University Health Centre for providing the infrastructure and resources that enabled this research, the RI-MUHC Molecular Imaging Core Platform for their valuable assistance with immunofluorescence imaging and EVE technologies for their expertise in multiplex ELISA.

## ABBREVIATIONS

ADAM17: A Disintegrin and Metalloproteinase Domain 17
ANXA3: Annexin A3
CBC: Complete blood counts
CCL: C-C motif chemokine ligand
CCR: C-C motif chemokine receptor
CDKN1B: Cyclin dependent kinase inhibitor 1B
CTLA4: Cytotoxic T-lymphocyte associated protein 4
cNLR: Circulating neutrophil-to-lymphocyte ratio
CKine: Chemokine
ELISA: Enzyme-linked immunosorbent assay
ENA-78: Epithelial neutrophil-activating peptide 78
FOXP3: Forkhead box P3
GEA: Gastroesophageal adenocarcinoma
GFP: Green fluorescent protein
GrA: Granzyme A
GrB: Granzyme B
H3Cit: Citrullinated histone H3
HAVCR2: Hepatitis A virus cellular receptor 2
ICOS: Inducible T-cell costimulator
IFN-γ: Interferon gamma
IL: Interleukin
MIP-3β: Macrophage inflammatory protein 3 beta
NET: Neutrophil extracellular trap
PB: Peripheral blood
PBMCs: Peripheral-blood mononuclear cells
PCNA: Proliferating cell nuclear antigen
PDGF: Platelet-derived growth factor
PD-L1: Programmed death ligand 1
PMA: Phorbol 12-myristate 13-acetate
RGCC: Regulator of cell cycle
RORC1/RORγ: Retinoic acid orphan receptor gamma
sFas: Soluble Fas
sFasL: Soluble Fas ligand
STAT: Signal transducers and activators of transcription
TNFRSF18: Tumour necrosis factor receptor subfamily member 18
Tp: Tumour periphery
TRAIL: Tumour necrosis factor-related apoptosis-inducing ligand
TRG: Tumour regression grade
VEGF-A: Vascular endothelial growth factor A.

