## Supplemental Material for "A Measurable Systemic Immune Phenotype Links Circulating Myeloid Dysfunction to Tumour Spatial Organization in Gastroesophageal Adenocarcinoma"

### SUPPLEMENTARY METHODS

#### S1.1 Patient Cohort and cNLR definitions

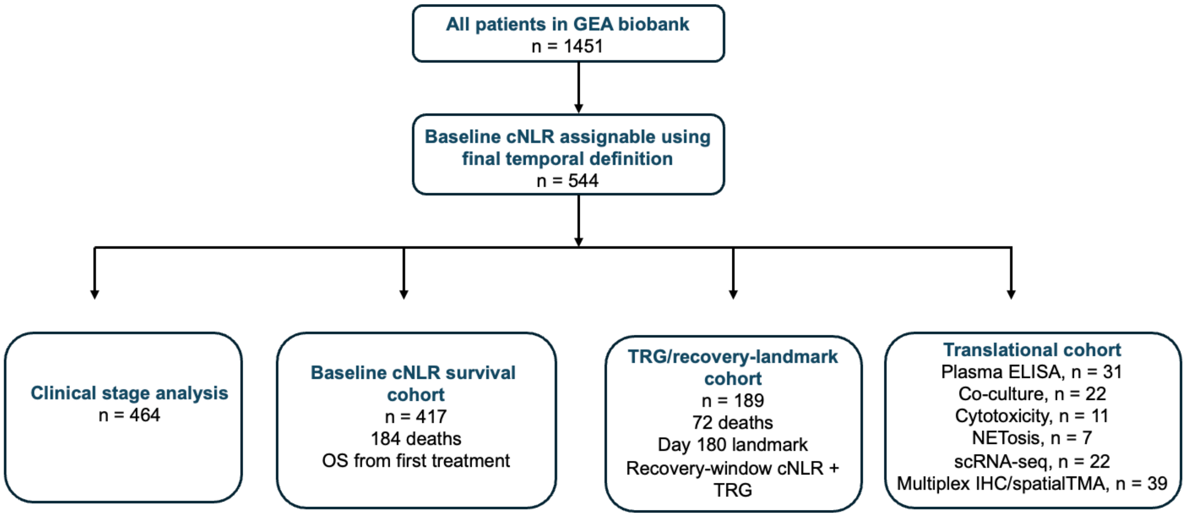


| Assay group | n | Plasma ELISA | Co-culture* | Cytotoxicity | NETosis | scRNA-seq | Spatial TMA |
|---|---|---|---|---|---|---|---|
| Plasma ELISA | 31 | — | | | | | |
| PBMC–neutrophil co-culture* | 22 | 1 | — | | | | |
| A549 cytotoxicity | 11 | 1 | 11 | — | | | |
| Spontaneous NETosis | 7 | 0 | 7 | 0 | — | | |
| scRNA-seq | 22 | 0 | 0 | 0 | 0 | — | |
| Spatial TMA | 39 | 7 | 0 | 0 | 0 | 0 | — |

**Supplementary Figure S1. Derivation of the clinical and translational cohorts and patient overlap across translational assays.** Derivation of the clinical and translational analysis cohorts and patient overlap across translational assays. The clinical analysis cohorts comprised 464 patients in the stage analysis, 417 patients in the baseline-survival cohort (184 deaths), and 189 patients in the recovery-window/TRG day-180 landmark cohort (72 deaths). The accompanying lower matrix reports the number of patients shared between each pair of translational assay group. Cohort sizes are shown in the n column. The scRNA-seq entry refers to the 22-patient resource, of which 7 pre-treatment tumour samples were evaluable for the neutrophil analysis. The spatial TMA comprised 39 patients, although one lacked an identifiable MRN for cross-cohort matching.

#### S1.1.1 Alternative cNLR summary measures.

To compare the prognostic information captured by different approaches to cNLR measurement, complementary single-measurement, window-summary, threshold-occupancy and longitudinal-change measures were evaluated. Pretreatment median cNLR was calculated as the median of valid daily pretreatment cNLR measurements. Pretreatment threshold occupancy was defined as the proportion of sampled pretreatment days with cNLR ≥4. Pretreatment change was expressed as the first-to-last pretreatment $\Delta\log_2$ cNLR. L90 cNLR was defined as described in the main Methods. Recovery-window cNLR was calculated as the median of valid daily postoperative measurements from postoperative days 60–180. Postoperative threshold occupancy was defined as the proportion of sampled postoperative days from days 60–180 with cNLR ≥4. Baseline-to-L90 change was expressed as $\Delta\log_2$ cNLR from baseline to L90. Record-wide median cNLR was calculated across all valid CBC measurements in the clinical record and was treated as an overall summary measure rather than a longitudinal trajectory. Threshold occupancy reflects the proportion of observed CBC days meeting the prespecified threshold and does not represent continuous time above threshold.

### S1.2 Detailed PMBC and neutrophil isolation

Peripheral blood mononuclear cells (PBMCs) and neutrophils were isolated from baseline peripheral blood using lymphocyte separation medium–based density-gradient centrifugation. Whole blood was diluted in calcium- and magnesium-free phosphate-buffered saline and carefully layered over 15 mL of lymphocyte separation medium

(Wisent Bioproducts; cat #305-010-CL). Samples were centrifuged at 800 × g for 30 minutes at room temperature without brake.

Following centrifugation, the PBMC layer at the plasma–lymphocyte separation medium interface was collected and washed twice in phosphate-buffered saline at 300 × g. Cell number and viability were assessed using a Vi-CELL automated cell counter (Beckman Coulter Life Sciences). Only PBMC samples with viability greater than 95% were used for downstream analyses.

The granulocyte- and erythrocyte-enriched lower fraction was collected, and erythrocytes were depleted by sedimentation in 6% dextran for 30 minutes. Residual erythrocytes were lysed using 1× BD Pharm Lyse Lysing Buffer (BD Biosciences; Cat. No. 555899) for 10 minutes at room temperature in the dark. Neutrophils were then washed twice in phosphate-buffered saline, recovered by centrifugation at 450 × g, and resuspended in the appropriate culture medium.

Neutrophil viability and purity were assessed using Trypan Blue and Methylene Blue staining, respectively. Only neutrophil preparations with a purity greater than 95% were used for downstream experiments. All samples were processed within 1–2 hours of blood collection.

#### S1.3 PMBC-neutrophil co-culture assay

Freshly isolated peripheral blood mononuclear cells (PBMCs) were cultured either alone or with autologous neutrophils to assess neutrophil-mediated effects on viable PBMC recovery. PBMCs were seeded at 1 × 10^6 cells per well in 24-well plates containing 1.5 mL of CTS OpTmizer T-Cell Expansion SFM medium (Gibco; Cat. No. A1048501).

PBMCs were stimulated with soluble anti-CD3 antibody at 1.0 μg/mL (Thermo Fisher Scientific; Cat. No. 16-0037-85) and anti-CD28 antibody at 2.0 μg/mL (Thermo Fisher Scientific; Cat. No. 16-0289-85). For coculture conditions, $5 \times 10^5$ autologous neutrophils were added at the time of stimulation, corresponding to a PBMC-to-neutrophil ratio of 2:1.

Cells were incubated for 5 days at 37°C in a humidified atmosphere containing 5% $CO_2$, without medium replacement or neutrophil removal. At the end of the culture period, viable PBMCs were quantified using a Vi-CELL automated cell counter.

Three technical replicate wells were performed for each patient and condition. The mean viable PBMC count from PBMC–neutrophil coculture wells was divided by the mean count from matched PBMC-alone wells to calculate relative viable PBMC expansion. A ratio below 1 indicated reduced viable PBMC expansion in the presence of neutrophils.

#### S1.4 A549-GFP cytotoxicity assay

PBMC-mediated cytotoxic activity was evaluated using GFP-expressing A549 adenocarcinoma cells as a standardized adherent tumour-cell target. A549-GFP cell line was generated by plasmid transfection from the original A549 cell line (Courtesy of Dr Sidong Huang, McGill University, Canada). A549-GFP cells were seeded at $5 \times 10^3$ cells per well in 96-well plates 24 hours before coculture in complete DMEM/F-12K medium (Wisent Bioproducts; cat #319-075-CL) containing 10% fetal bovine serum (Wisent Bioproducts; cat #090150) and 1% penicillin–streptomycin (Wisent Bioproducts; cat #450-201-EL) and incubated at 37°C.

PBMCs recovered from the preceding PBMC-alone or PBMC–neutrophil coculture conditions were added at $2.5 \times 10^5$ cells per well in 200 μL of T-cell expansion medium, corresponding to an effector-to-target ratio of 50:1. Cocultures were incubated for 24 hours at 37°C in a humidified atmosphere containing 5% $CO_2$.

Following incubation, wells were gently washed with phosphate-buffered saline to remove PBMCs and nonadherent A549-GFP cells. One image was acquired from the central field of each well using an EVOS imaging system under identical exposure settings. Residual adherent GFP-positive tumour-cell area was quantified in ImageJ v2.1. Lower GFP-positive area was interpreted as greater PBMC-mediated cytotoxic activity.

Six technical replicate wells were performed for each patient and condition and averaged to generate one patient-level value. Image acquisition and quantification were performed blinded to cNLR group.

#### S1.5 NET formation and immunofluorescence

Neutrophil extracellular trap formation was assessed in freshly isolated neutrophils from 7 patients stratified according to baseline cNLR (n=4 low; 3 high NLR). Neutrophils were seeded at $1 \times 10^6$ cells per well in 24-well plates containing 1 mL of serum-free RPMI medium (Wisent Bioproducts; cat #350-000-CL) and incubated for 4 hours at 37°C in a humidified atmosphere containing 5% $CO_2$ without stimulation. For positive controls, neutrophils were treated with 250 nM phorbol 12-myristate 13-acetate (MedChem Express; Cat. No. HY-18739-10mg) under identical culture conditions. 3 wells were plated per patient due to cell availability, one for positive control, one for SYTOX staining and one for immunofluorescence staining.

At the end of the incubation period, SYTOX Green was added at a final concentration of 250 nM and incubated for 5 minutes at 37°C to label extracellular DNA. NET formation was visualized using an EVOS imaging system. Three to six randomly selected fields per well were acquired per patient under identical exposure settings, and characteristic web-like extracellular DNA structures were used as a qualitative indicator of NETosis. Fields with no cells/NETs were excluded.

For immunofluorescence confirmation and quantification, cells were fixed overnight in 4% paraformaldehyde. The following day, paraformaldehyde was removed and cells were permeabilized with 0.1% Triton X-100 (Thermo Fisher Scientific; cat #A16046.AE) in phosphate-buffered saline for 60 seconds, followed by two 5-minute washes in phosphate-buffered saline. Samples were then blocked for 1 hour at room temperature in Tris-buffered saline containing 0.1% Tween-20, 1% bovine serum albumin, 1% fetal bovine serum, and 0.01% Triton X-100.

Cells were incubated for 1 hour at room temperature with rabbit anti-human citrullinated histone H3 antibody at 1:100 dilution (Novus Biologicals; Cat. No. NBP3-11316) and anti-human CD66b-APC antibody at 1:50 dilution (clone REA306; Miltenyi Biotec; Cat. No. 130-135-357). Following two 5-minute washes in phosphate-buffered saline, cells were incubated for 1 hour at room temperature with goat anti-rabbit IgG Alexa Fluor 568 secondary antibody at 1:500 dilution (Thermo Fisher Scientific; Cat. No. A-11011) and DAPI at 1:2000 dilution (Thermo Fisher Scientific; Cat. No. D1306). Samples were then washed twice in phosphate-buffered saline and stored in 1 mL phosphate-buffered saline at 4°C in the dark until imaging.

Fluorescence images were acquired using a Zeiss LSM 880 confocal microscope with a 20× objective. Four randomly selected fields were captured per condition using identical acquisition settings. A negative control (DAPI + secondary antibodies only) was included to assess tissue autofluorescence and non-specific antibody binding and was used to establish image acquisition parameters and fluorescence intensity thresholds for subsequent ImageJ v2.1 analysis. NETs were identified as H3Cit-positive structures associated with CD66b-positive neutrophils. Quantification was performed in ImageJ v2.1 by calculating the H3Cit-positive area divided by the CD66b-positive area for each field. Image acquisition and quantification were blinded to cNLR group.

For each patient and condition, four fields were retained as separate technical measurements. Fields were not treated as independent biological replicates; they were analyzed using a linear mixed-effects model with NLR group as a fixed effect and patient identity as a random intercept to account for field clustering within patient. Identical image-processing parameters were applied across samples. Higher H3Cit-positive area normalized to CD66b-positive area was interpreted as greater NET formation.

#### S1.6 Plasma cytokine and chemokine profile

Plasma samples collected at diagnosis from treatment-naïve GEA patients were analysed for soluble immune mediators. Patients were stratified according to baseline circulating neutrophil-to-lymphocyte ratio into low-cNLR (<4; n = 17) and high-cNLR (≥4; n = 14) groups.

For each patient, 200 µL of plasma was analysed by EVE Technologies using the Human Cytokine/Chemokine 96-Plex Discovery Assay according to the manufacturer's protocol. Samples were assayed in technical duplicate, and duplicate measurements were averaged to generate a single concentration for each analyte and patient. The complete analyte list is provided in **Supplementary Table S2**.

To assess coordinated biological processes, predefined composite scores were generated from functionally related analytes. Individual analyte concentrations were standardized across the study cohort by calculating z-scores. For each patient, the composite score was calculated as the arithmetic mean of the standardized values for all analytes included in the corresponding biological signature, thereby assigning equal weight to each component. The analytes included in each composite score are listed in **Supplementary Table S2**.

Patients with a missing value for any analyte included in a given composite were excluded from that specific analysis. Composite-score comparisons between low- and high-cNLR groups were performed using Welch's two-sample t test, with the Mann–Whitney U test used as a nonparametric sensitivity analysis. Effect sizes were summarized using Cohen's d. All tests were two-sided, and $P < .05$ was considered statistically significant.

#### S1.7 scRNA-seq tissue processing and library preparation

Fresh tumour specimens were trimmed to remove necrotic areas and enzymatically dissociated in Advanced DMEM/F12 containing collagenase type III (10 mg; Worthington) and hyaluronidase (500 U; Sigma) using C-tubes on a gentleMACS Octo Dissociator (Miltenyi Biotec). The resulting suspension was processed in phosphate-buffered saline containing 1 mM dithiothreitol, filtered through a 100-μm strainer, and centrifuged at 500 × g for 5 minutes at 4°C.

Cells were further incubated with 0.25% trypsin–EDTA for 5 minutes at 37°C. Enzymatic activity was quenched with 10% fetal bovine serum, and the suspension was pelleted and treated with Dispase (2.5 U) and DNase (10 μg) for 5 minutes at 37°C. Following quenching in phosphate-buffered saline, the suspension was passed through a 40-μm strainer and centrifuged. Residual erythrocytes were removed using ACK lysis buffer (Thermo Fisher Scientific; cat # A1049201) for 5 minutes at room temperature, followed by two washes in phosphate-buffered saline containing 2% fetal bovine serum.

Single-cell suspensions were assessed for viability, debris, erythrocyte contamination, and cell concentration before capture. Preparations were required to meet predefined quality thresholds of at least 70% viability, no more than 10% erythrocytes, and no more than 30% debris. Viability was assessed using Calcein-AM and Ethidium Homodimer-1 staining from the LIVE/DEAD kit (Thermo Fisher Scientific; cat #L3224) and quantified by fluorescence microscopy. Erythrocyte contamination was evaluated using DRAQ5 nuclear staining and bright-field morphology. Cell concentration was determined by hemocytometer counting with standard dilution correction.

Eligible suspensions were processed using the 10x Genomics Chromium platform with Chromium Next GEM Single Cell 3′ v3.1 chemistry.

Library concentrations were quantified using KAPA quantitative PCR (Roche). For MGI sequencing, 10x libraries were converted using the MGIEasy Universal Library Conversion Kit, circularized, and amplified into DNA nanoballs before sequencing on DNBSEQ-G400 instruments using paired-end 100- or 150-base-pair reads with App-A chemistry. A subset of libraries was sequenced on Illumina NovaSeq 6000 or HiSeq 4000 instruments using standard pooling and loading procedures.

Raw MGI data were demultiplexed using fastq-multx and fgbio, whereas Illumina data were processed using bcl2fastq. FASTQ files generated from multiple sequencing runs were merged by library before downstream processing.

### S1.8 scRNA-seq computational analysis

#### S1.8.1 Read processing and count-matrix generation

Sequencing reads were trimmed using cutadapt version 3.2 and pseudo-aligned to the GRCh38 transcriptome using Ensembl release 96 and kallisto version 0.46.2. Gene-level count matrices were generated using bustools version 0.40.0. Initial quality control, normalization, dimensionality reduction, and clustering were performed using Scanpy version 1.7.1. Subsequent statistical and pathway analyses were performed in R version 4.5.1 using RStudio version 2024.09.1.

#### S1.8.2 Cell-level quality control

Cells with fewer than 1,000 unique molecular identifiers, fewer than 500 detected genes, or more than 10% mitochondrial reads were excluded. Doublets were identified using Scrublet (0.2.4) and excluded at a score threshold greater than 0.2. Only protein-coding genes were retained for downstream analysis. Following quality control, a total of 73,844 cells from 26 samples from 22 patients were retained (n=13 low-, 9 high cNLR).

#### S1.8.3 Normalization, integration, clustering, and visualization

Count matrices were normalized, log-transformed, and scaled after identification of 4,000 highly variable genes. Principal-component analysis was performed using 50 components. Samples were integrated using Harmony, after which a k-nearest-neighbour graph was constructed using $k = 15$. Leiden clustering was performed on the integrated graph, and clusters were visualized using uniform manifold approximation and projection initialized with partition-based graph abstraction coordinates.

#### S1.8.4 Cell-type annotation

Initial cell identities were assigned using a previously published pipeline developed by Strasser et al. Expression profiles were mapped to the Human Cell Landscape reference after library-size normalization and log transformation. For each cell, Pearson correlation was calculated against reference profiles, and the label with the maximum correlation was assigned. Cells with a maximum correlation below 0.3 were classified as unknown.

Cluster-level annotations were subsequently reviewed and refined using canonical marker-gene expression. Marker selections were curated from the Cell Ontology and established antibody and gene-expression resources.

Given the limited neutrophil representation across the cohort, subsequent neutrophil-specific analysis was restricted to 7 pre-treatment tumour samples (n = 3 low cNLR, 4 high cNLR).

#### S1.8.5 Functional gene-signature analysis

Functional gene-signature analysis was performed on neutrophils from the 7 pre-treatment tumour samples described above at the single cell level. Customized functional gene signatures were curated to represent neutrophil -associated biological programmes described in the literature.

Signature activity was quantified using UCell on normalized expression matrices from the relevant neutrophil populations. UCell generated rank-based signature scores for individual cells. For between-group analyses, UCell scores were summarized at the patient level or modelled while accounting for clustering of cells within patients.

#### S1.8.6 Differential expression of selected genes

Differential expression of pre-selected genes between high- and low-cNLR groups was assessed separately in neutrophils (n = 7 patients) and T cells at the single-cell level using the FindMarkers function in Seurat (5.4.0), with the Wilcoxon rank-sum test (**Fig. 4C,E**). Genes with a nominal $P < 0.05$ were visualised, with significance indicated according to Benjaminin Hochberg adjusted P value.

#### S1.8.7 Patient level pathway-enrichment analysis

Sample- and cell-type-specific pseudobulk profiles were generated separately for different cell compartments. Reactome and Hallmark collection pathway activity was evaluated using gene set variation analysis with the GSVA R package (2.4.4), using sample-level pseudobulk expression profiles as input. For patients represented by multiple samples, sample-level GSVA scores were averaged for each pathway to obtain a single score per patient. Immune-system-related Reactome pathways were compared between low- and high-cNLR groups **(Fig. 4A,B)**, while Hallmark pathway analyses were performed across individual cell compartments **(Fig. 4F)**. Patient-level pathway scores were compared between cNLR groups using a two-sided Wilcoxon rank-sum test, with *P* values adjusted using the Benjamini–Hochberg method.

#### S1.8.8 Softwares

The main software versions were cutadapt 3.2, kallisto 0.46.2, bustools 0.40.0, Scanpy 1.7.1, Scrublet 0.2.3, Harmony 1.2.4, Seurat 5.4.0, GSVA 2.4.4.

#### S1.9 Multiplex immunostaining and assay optimization

Endoscopic biopsy or surgical resection specimens containing tumour and, where available, histologically normal adjacent tissue were used for tissue microarray construction. Tissue regions were selected by a pathologist following review of hematoxylin and eosin–stained sections. Tumour core was defined as central malignant tissue, tumour periphery as the tumour–stroma interface, and adjacent non-malignant tissue as histologically normal tissue without tumour involvement. Initial chromogenic immunohistochemistry for CD3 and neutrophil elastase was performed to quantify lymphocytes and neutrophils, respectively, and to calculate tissue neutrophil-to-lymphocyte ratio.

Multiplex immunofluorescence staining was performed by the Molecular and Cellular Immunology Core at the BC Cancer Victoria Deeley Research Centre using the Akoya OPAL tyramide signal amplification system on an IntelliPATH FLX automated staining platform (Akoya Biosciences). Formalin-fixed paraffin-embedded tissue sections were stained according to institutional standard operating procedures and manufacturer recommendations.

##### S1.9.1 Tissue preparation and antigen retrieval

Formalin-fixed paraffin-embedded slides were baked at 37°C overnight or, when required, at 58°C for 30 minutes, followed by manual deparaffinization and rehydration. Slides were post-fixed in 10% neutral buffered formalin (Sigma-Aldrich; Cat. No. HT501128-4L) for 20 minutes and rinsed thoroughly in distilled water.

Heat-induced epitope retrieval was performed using Biocare Nuclear Decloaker solution (Cat. No. BC-CB911 M) in a decloaking chamber at 110°C for 15 minutes, followed by 100°C and passive cooling.

#### S1.9.2 Sequential multiplex staining

Staining was performed in iterative cycles on the IntelliPATH FLX platform (**Supplementary Table S3**). Following each staining round, slides were washed in Tris-buffered saline (Biocare; Cat. No. BC-TWB945). Endogenous peroxidase activity was quenched using Peroxidazed-1 (Biocare; Cat. No. BC-PX968), followed by blocking with Background Sniper (Biocare; Cat. No. BC-BS966) or Background Terminator (Biocare; Cat. No. BC-BT967), depending on the primary antibody.

Primary antibodies were applied sequentially according to the staining panel provided in Supplementary Table S3. Species-specific horseradish peroxidase–conjugated secondary polymers were then applied (Biocare; Cat. No. BC-RHRP520). Signal amplification was performed using OPAL fluorophores diluted in amplification buffer and incubated for 10 minutes at room temperature in the dark. Fluorophores were prepared immediately before use because of their light sensitivity.

Between staining cycles, antibody–fluorophore complexes were stripped by antigen retrieval using either AR6 buffer (Akoya Biosciences; Cat. No. AR600250ML) or AR9 buffer (Akoya Biosciences; Cat. No. AR900250ML). Retrieval was performed using either a decloaking chamber or a microwave-based DAVE OPAL method, according to the optimized requirements of the antibody panel. Slides were cooled, washed extensively in water, and returned to the staining platform for the subsequent cycle.

#### S1.9.3 Final staining and counterstaining

In the final staining cycle, OPAL 780 detection (Akoya Biosciences; Cat. No. FP1501001KT) was performed manually because of its high light sensitivity. Sections were incubated with the corresponding primary antibody for 1 hour at room temperature, followed by OPAL 780 detection.

Nuclei were counterstained with DAPI for 5 minutes (Akoya Biosciences; Cat. No. FP1490). Slides were then washed and mounted using ProLong Diamond Antifade Mountant (Thermo Fisher Scientific; Cat. No. P3696).

#### S1.9.4 Antibody panel and assay controls

The multiplex panel included markers for granulocytes (CD66b), T-cell subsets (CD4, CD8, and FOXP3), CD68-positive myeloid cells, immune-checkpoint expression (PD-L1), and epithelial cells (pan-cytokeratin), together with DAPI nuclear counterstaining. Antibody clones, concentrations, retrieval conditions, and fluorophore assignments are provided in Supplementary Table S3.

Control tissues included tonsil for CD4, CD8, FOXP3, and CD66b; spleen for PD-L1 and CD68; and tumour epithelium for pan-cytokeratin. Expected staining patterns included membranous CD4 and CD8 expression in T cells, nuclear FOXP3 expression in regulatory T cells, cytoplasmic CD68 expression in myeloid cells, membranous PD-L1 expression in immune and tumour compartments, and cytoplasmic pan-cytokeratin expression in epithelial cells.

Single-marker controls were used for spectral-library development and assessment of staining specificity. Autofluorescence controls were used during spectral unmixing.

#### S1.9.5 Image acquisition

Slides were imaged using a Vectra Polaris imaging system (Akoya Biosciences) at 20× magnification. Multispectral image sets were acquired under standardized imaging conditions and processed using inForm software v10.1.2 (Akoya Biosciences).

### S1.10 Tissue segmentation and cellular phenotyping

#### S1.10.1 Tissue and cellular segmentation

A supervised machine-learning algorithm was developed using a training set of 15 representative images. The trained model was used to segment tissue into epithelial, stromal, and other regions and to perform individual-cell segmentation.

DAPI, pan-cytokeratin, and autofluorescence channels were used to define nuclear boundaries, cellular morphology, and tissue compartments. Tissue segmentation and cell segmentation were reviewed for technical adequacy before quantitative analysis.

#### S1.10.2 Cellular phenotyping

Eleven phenotyping models were developed to classify mutually exclusive cellular phenotypes while preserving the X–Y coordinates of each cell. Phenotype definitions and marker combinations are provided in **Supplementary Table S4**.

Quantitative phenotype outputs were generated using PhenoptrReports v0.3.3 (Akoya Biosciences), which integrated results from the separate phenotyping models to calculate cell counts, cell densities, and spatial distributions for each defined phenotype within epithelial and stromal compartments.

### Measurements from multiple image fields or tissue cores within the same anatomical compartment were summarized at the patient level for downstream analyses.**S1.11 Spatial Analysis implementation**

#### S1.11.1 Image fields, anatomical compartments and patient-level aggregation

Cell-level multiplex immunofluorescence data were analysed using phenotype assignments and X–Y centroid coordinates generated by the image-analysis pipeline. Image fields were linked to the clinical database through the locked TMA core map and assigned to harmonized anatomical categories, including tumour core, tumour periphery, normal stomach, normal esophagus and premalignant field. Premalignant-field tissue comprised non-invasive Barrett's-lineage cores annotated as Barrett's, Barrett's with dysplasia, or dysplasia.

Analyses were initially performed at the image-field level. When multiple evaluable fields represented the same patient and anatomical compartment, field-level measurements were summarized using the median to generate one patient-level value.

#### S1.11.2 Cell populations and estimated abundance

The principal populations evaluated were $CD8^+$ T cells, $CD66b^+$ neutrophils, $CD68^+$ macrophages, $PD\text{-}L1^+CD66b^+$ neutrophils and $PD\text{-}L1^+CD68^+$ macrophages. A combined $PD\text{-}L1^+$ myeloid population comprised cells meeting either $PD\text{-}L1^+CD66b^+$ or $PD\text{-}L1^+CD68^+$ criteria, with dual-positive cells counted once.

Phenotype-positive cell counts were calculated separately for each image field. Analysed field area was estimated from the rectangular coordinate extent. Coordinates were converted to micrometres using a calibration of 0.5 μm per pixel, and area was converted to mm². Estimated phenotype density was calculated as the phenotype-positive cell count divided by estimated analysed field area. Because the denominator was derived from the cellular-coordinate extent rather than an exact epithelial or stromal segmentation mask, these values were interpreted as estimated field densities.

#### S1.11.3 Nearest-neighbour proximity and clustering

Spatial relationships between $CD8^+$ T cells and myeloid populations were quantified using Euclidean centre-to-centre distances. For each $CD8^+$ cell, the distance to the nearest specified myeloid cell was calculated. Coincident zero-distance matches attributable to the same or dual-labelled segmented cell were excluded before identifying the nearest distinct neighbour.

Field-level summaries included the median nearest-neighbour distance and the proportions of $CD8^+$ T cells within 25, 50 and 100 μm of the specified myeloid population.

Within-population clustering was evaluated for $CD8^+$, $CD66b^+$, $CD68^+$, PD-$L1^+CD66b^+$, PD-$L1^+CD68^+$ and combined PD-$L1^+$ myeloid populations. For populations containing at least two evaluable cells, observed nearest-neighbour spacing was compared with that expected under complete spatial randomness. Clustering scores were oriented such that positive values indicated shorter-than-expected spacing and greater clustering, whereas negative values indicated relative dispersion.

#### S1.11.4 Cross-compartment and pooled tumour measures

The principal premalignant-field spatial gradient used in **Fig. 5H** was calculated by subtracting the patient-level premalignant-field median CD8-to-PD-$L1^+CD68^+$ nearest-neighbour distance from the corresponding tumour-periphery value. Positive values therefore indicated greater CD8-to-PD-L1$^{+}$CD68$^{+}$ separation at the tumour periphery relative to the premalignant field.

For pooled malignant-compartment abundance analyses, tumour CD66b$^{+}$ abundance was summarized as the patient-level median across tumour-core and tumour-peripheral CD66b$^{+}$ density measurements

#### S1.11.5 Statistical analysis

Spatial metrics were analysed as continuous variables unless otherwise specified. Paired anatomical comparisons used the Wilcoxon signed-rank test, independent-group comparisons used the Wilcoxon rank-sum test, and associations with continuous cNLR measures were assessed using Spearman rank correlation. For categorical cNLR analyses, the prespecified threshold of $<4$ versus $\geq 4$ was used.

Associations with recurrence-free survival (RFS) were evaluated using log-rank testing and univariable Cox proportional-hazards models. Documented recurrence was treated as the event; patients without documented recurrence, including those who died without prior recurrence, were censored. Continuous spatial variables were standardized before Cox modelling so that hazard ratios represented a one-standard-deviation increase. For visualization of the continuous tumour-periphery PD-L1$^{+}$ myeloid clustering association shown in **Fig. 5F**, model-estimated RFS curves were displayed at the first and third quartiles of the clustering score rather than using a dichotomized cutoff.

As a sensitivity analysis, the premalignant-field definition was broadened to include normal stomach; associations involving premalignant-field spatial measures were attenuated under this broader definition. All spatial analyses were exploratory and two-sided. Analysis-specific eligibility, missing-data handling and evaluable sample sizes are reported with the corresponding results and figure panels.

### SUPPLEMENTARY TABLES

#### Supplementary Table S1: Clinical characteristics of the overall baseline cNLR cohort and assay-specific translational cohorts.

| Characteristic | Overall baseline cNLR cohort | ELISA | Co-culture | Cytotoxicity | NETosis | scRNA seq | scRNA-seq neutrophil cohort | Spatial TMA |
|---|---|---|---|---|---|---|---|---|
| Patients, n | 464 | 31 | 22 | 11 | 7 | **22** | **7** | 39 |
| Age at diagnosis, median (IQR) | 67.7 (60.2–74.2) | 67.0 (60.8–76.0) | 70.0 (64.5–75.2) | 73.0 (68.5–75.5) | 71.0 (60.0–75.0) | 68.4 (63.1-73.0) | 66.0 (61.0–70.0) | 66.0 (59.0–70.0) |
| Male sex, n/N (%) | 358/463 (77.3%) | 25/31 (80.6%) | 20/22 (90.9%) | 10/11 (90.9%) | 7/7 (100.0%) | 14/22 (63.6%) | 6/7 (85.7%) | 33/39 (84.6%) |
| **Baseline cNLR, median (IQR)** | 3.43 (2.37–5.05) | 3.38 (2.31–5.24); n=31 | - | - | - | - | 4.01 (3.55–4.26); n=6 | 2.83 (2.18–5.21); n= 39 |
| **cNLR ≥4, n/N (%)** | 142/464 (30.6%) | 14/31 (45.2%) | 9/22(40.9%) | 4/11 (36.4%) | 3/7 (42.9%) | 9/22 (40.9%) | 4/7 (57.1%) | 11/39 (28.2%) |
| Stage I, n/N (%) | 43/464 (9.3%) | 7/30 (23.3%) | 1/22 (4.5%) | 1/11 (9.1%) | 0/7 (0.0%) | 2/22 (9.1%) | 0/7 (0.0%) | 4/39 (10.3%) |
| Stage II, n/N (%) | 75/464 (16.2%) | 8/30 (26.7%) | 11/22 (50.0%) | 5/11 (45.5%) | 5/7 (71.4%) | 5/22 (22.7%) | 0/7 (0.0%) | 12/39 (30.8%) |
| Stage III, n/N (%) | 249/464 (53.7%) | 11/30 (36.7%) | 5/22 (22.7%) | 3/11 (27.3%) | 1/7 (14.3%) | 12/22 (54.5%) | 7/7 (100.0%) | 19/39 (48.7%) |
| Stage IV, n/N (%) | 97/464 (20.9%) | 4/30 (13.3%) | 5/22 (22.7%) | 2/11 (18.2%) | 1/7 (14.3%) | 3/22 (13.6%) | 0/7 (0.0%) | 4/39 (10.3%) |
| Esophagus, n/N (%) | — | 9/31 (29.0%) | 8/22 (36.4%) | 3/11 (27.3%) | 3/7 (42.9%) | 13/22 (59.1%) | 1/7 (14.3%) | 7/11 (63.6%) |
| GEJ, n/N (%) | — | 15/31 (48.4%) | 6/22 (27.3%) | 4/11 (36.4%) | 2/7 (28.6%) | 8/22 (36.4%) | 6/7 (85.7%) | 4/11 (36.4%) |
| Gastric, n/N (%) | — | 7/31 (22.6%) | 8/22 (36.4%) | 4/11 (36.4%) | 2/7 (28.6%) | 1/22 (4.5%) | 0/7 (0.0%) | 0/11 (0.0%) |
| Neoadjuvant treatment, Y / N | 262 / 116 / 86 | 16 Y / 12 N; 3 N/A | 14 Y / 4 N; 4 N/A | 7 Y / 2 N; 2 N/A | 4 Y / 2 N; 1 N/A | 14 Y/ 7 N; 1 N/A | 7 Y / 0 N | 30 Y / 9 N |

Values are median (IQR) or n/N (%), unless otherwise indicated. Denominators reflect available clinical annotation within each assay-specific cohort. For some translational cohort patients, only categorical baseline cNLR classification (low <4 vs high ≥4) was available, whereas a numerical baseline cNLR value could not be recovered; therefore, denominators may differ between the continuous baseline cNLR and cNLR ≥4 rows.

**Supplementary Table S2: Complete Panel of Soluble Immune Mediators Quantified in Patient Plasma in 96-Plex Discovery Assay.** Plasma samples from treatment-naïve GEA patients were analyzed using the Human Cytokine/Chemokine 96-Plex Discovery Assay from Eve Technologies. Analytes include cytokines, chemokines, growth factors, angiogenic factors, hematopoietic factors, cell death mediators and immune mediators. Analytes were grouped according to their predominant biological function for ease of interpretation. Markers in italics were not excluded in composite analysis due to confounding pleotropic functions.

| Functional Category | Analytes |
|---|---|
| Cell Death | sFas, TRAIL, Granzyme A, Granzyme B, Perforin |
| Angiogenesis | VEGF-A, PDGF-AA, PDGF-AB/BB, ENA-78, *EGF, FGF-2* |
| Inflammatory Hematopoiesis | G-CSF, GM-CSF, IL-3, M-CSF, SCF, FLT-3L, TPO, LIF |
| Immunosuppression | IL-10, IL-1RA, IL-35, APRIL, BAFF, sCD137, sCD40L, TSLP |
| Innate Inflammation | I-309, MIP3β, IL-18, IL-28A, IFNω, 6CKine, sFasL |
| Adaptive Immune Activation | IL-2, IL-4, IL-5, IL-7, IL-9, IL-11, IL-12p40, IL-12p70, IL-13, IL-15, IL-16, IL-17A, IL-17E/IL-25, IL-17F, IL-20, IL-21, IL-22, IL-23, IL-24, IL-27, IL-31, IL-33, IL-34 |
| Interferon Signaling | IFN-α2, IFNβ, IFNγ, IL-29 |
| Myeloid-Related | GCP-2, GROα, CXCL16, MCP-1, MCP-2, MCP-3, MCP4, MIP1α, MIP1β, MIP1δ, MPIF-1, HMGB1, IL-1α, IL-1β, TNFα, TNFβ, IL-8 |
| Lymphocyte-Related | BCA-1, CCL28, CTACK, Eotaxin, Eotaxin-2, Eotaxin-3, Frackaltine, Lymphotactin, MDC, MIG/CXCL9, MIP3α, RANTES, SDF-1, TARC, IP-10, I-TAC |

**Abbreviations:** APRIL, a proliferation-inducing ligand; BAFF, B-cell activating factor; EGF, epidermal growth factor; FGF, fibroblast growth factor; FLT-3L, FMS-like tyrosine kinase 3 ligand; G-CSF, granulocyte colony-stimulating factor; GM-CSF, granulocyte-macrophage colony-stimulating factor; HMGB1, high-mobility group box 1; IFN, interferon; IL, interleukin; LIF, leukemia inhibitory factor; MCP, monocyte chemoattractant protein; M-CSF, macrophage colony-stimulating factor; MDC, macrophage-derived chemokine; MIP, macrophage inflammatory protein; PDGF, platelet-derived growth factor; SCF, stem cell factor; SDF, stromal cell-derived factor; TGF, transforming growth factor; TNF, tumour necrosis factor; TPO, thrombopoietin; TRAIL, tumour necrosis factor–related apoptosis-inducing ligand; TSLP, thymic stromal lymphopoietin; VEGF, vascular endothelial growth factor.

**Supplementary Table S3: Summary of Antibodies Used for 8-plex Immunostaining of Tumour Microarrays.** The antibodies are listed in the order in which they were sequentially stained in. All the fluors listed here are from Akoya Biosciences. Abbreviations: Rb = rabbit; Ms = mouse; TSA-DIG = Tyramide Signal Amplification-Digoxigenin

| Round | $1^0$ Ab | Host | Clone | Cat # | AR Before | Block | Dilution | Polymer | OPAL Fluor | Cat# Fluor |
|---|---|---|---|---|---|---|---|---|---|---|
| **1** | CD66b | Rb | EPR25354-2 | ab300122 | Nuclear | Sniper 10 | 1/20000 | M2 Rb HRP | 620 | FP1495 001KT |
| **2** | CD8 | Ms | C8/144B | ab17147 | AR6 | Sniper 10 | 1/600 | M2Ms-HRP | 570 | FP1488 001KT |
| **3** | CD4 | Rb | EPR6855 | ab133616 | AR6 | Sniper 10 | 1/500 | M2 Rb HRP | 480 | FP1500 001KT |
| **4** | CD68 | Rb | D4B9C | 76437 | AR6 | Sniper 10 | 1/500 | M2Rb-HRP | 690 | FP1497 001KT |
| **5** | FoxP3 | Ms | 236A/E7 | 501129542 | AR6 | Sniper 10 | 1/100 | M2Ms-HRP | 540 | FP1494 001KT |
| **6** | PD-L1 | Rb | SP142 | ab228462/ M4422 | AR9 | Sniper 5 | 1/100 | M2 Rb HRP | 520 | FP1487 001KT |
| **7** | PanCK + | Ms | AE1/AE3+5D3 | CM162 | AR6 | Sniper 10 | 1/100 | M2 Ms HRP | TSA-DIG | N/A |
| **8** | 780 | N/A | N/A | FP1501001 KT | AR9 | N/A | 1/25 | N/A | N/A | N/A |

**Supplementary Table S4: Phenotype Models Used for Human Tumour Microarray Analysis.** The 11 phenotypes as described below were used for the analysis of 8-plex immunostaining of the human tumour microarray. Red denotes negativity for the marker, while green denotes positivity for the respective marker. Abbreviations: Tregs = Regulatory T cells; PanCK = Pan Cytokeratin

| | **CD4** | **CD8** | **CD66b** | **CD68** | **FOXP3** | **PD-L1** | **PanCK** |
|---|---|---|---|---|---|---|---|
| **Cancer cells** | Red | Red | Red | Red | Red | Red | Green |
| **T Helper cells** | Green | Red | Red | Red | Red | Red | Red |
| **CD8+ T cells** | Red | Green | Red | Red | Red | Red | Red |
| **CD68+ myeloid cells** | Red | Red | Red | Green | Red | Red | Red |
| **Tregs** | Red | Red | Red | Red | Green | Red | Red |
| **Neutrophils** | Red | Red | Green | Red | Red | Red | Red |
| **PDL1+ CD66b+ neutrophils** | Red | Red | Green | Red | Red | Green | Red |
| **Tumour PD-L1** | Red | Red | Red | Red | Red | Green | Green |
| **PD-L1+ CD4+ T cells** | Green | Red | Red | Red | Red | Green | Red |
| **PD-L1+ CD8+ T cells** | Red | Green | Red | Red | Red | Green | Red |
| **PD-L1+ CD68+ myeloid cells** | Red | Red | Red | Green | Red | Green | Red |

**Supplementary Table S5. Multivariable Cox proportional-hazards model for overall survival according to baseline cNLR in the locked baseline-survival cohort.** The number of patients is the cohort is 417, with 184 recorded deaths.

| Variable | Adjusted HR | 95% CI | P |
|---|---|---|---|
| Baseline cNLR, per doubling | 1.30 | 1.10–1.54 | .002 |
| Clinical stage II vs I | 1.68 | 0.84–3.36 | .141 |
| Clinical stage III vs I | 1.43 | 0.78–2.65 | .250 |
| Clinical stage IV vs I | 3.18 | 1.67–6.08 | <.001 |
| Age, per 10 years | 1.04 | 0.90–1.20 | .566 |
| Male vs female | 1.14 | 0.78–1.67 | .488 |
| Esophageal/GEJ cohort vs gastric cohort | 1.15 | 0.79–1.68 | .457 |

**Supplementary Table S6. Cox regression analysis of recovery-window cNLR, pathological tumour regression grade, and overall survival in the day-180 landmark cohort.**

| Model | Variable | Adjusted HR | 95% CI | P |
|---|---|---|---|---|
| **Categorical model** | Poor TRG vs good TRG | 2.14 | 1.20–3.82 | .010 |
| | Recovery-window cNLR ≥4 vs <4 | 2.57 | 1.53–4.29 | <.001 |
| **Continuous model** | Recovery-window cNLR, per doubling | 1.43 | 1.11–1.85 | .006 |

Note: N=189, 72 deaths; models stratified by primary tumour site; categorical model mutually adjusted TRG and cNLR.
